# Environmental variability can promote parasite diversity within hosts and transmission among hosts

**DOI:** 10.1101/2023.06.05.543787

**Authors:** Madeline Jarvis-Cross, Martin Krkosek

## Abstract

While the mechanisms that govern disease emergence and spread among hosts are increasingly well-described, the mechanisms that promote parasite diversity within-hosts, affecting host outcomes and spillover potential, have been comparatively understudied. Furthermore, while attention has been paid to the effects of increasing temperatures on disease systems, the effects of environmental variability have been left underexplored, despite rising climatic variability. To investigate the impacts of environmental variability on parasite diversity within-hosts, we analyzed a model of within-host population dynamics wherein two parasites indirectly compete through the host’s immune response. We simulated the model under constant, demographically stochastic, environmentally stochastic, and demographically and environmentally stochastic conditions, and analysed the viability and longevity of non-equilibrium parasite co-occurrence. We found that environmental stochasticity increased the viability and longevity of parasite co-occurrence, suggesting that thermal variability arising from climatic change and as a physiological trait may promote parasite diversity within ectotherms and help explain bats’ propensity to support diverse communities of parasites. Further, we found that under certain conditions, the transmissibility of co-occurring parasites can surpass the transmissibility of single parasites, suggesting that thermal variability may increase the transmissibility of co-occurring parasites.

## Introduction

It has become clear that climate and land-use change can facilitate novel species interactions and increase contact rates between wild populations and humans, resulting in devastating epidemics and enabling spillover and disease emergence events among wild and human hosts alike (Daszak et al., 2000, 2001; Plowright et al., 2017). There is strong theoretical and empirical evidence that within-host parasite diversity influences disease outcomes in individual hosts (Bromenshenk et al., 2010; Johnson & Hoverman, 2012), transmissibility among hosts (Jolles et al., 2008; Susi et al., 2015), and community structure and functioning (Lafferty et al., 2006). However, the mechanisms that promote within-host parasite diversity have been understudied.

Ecological theory predicts a relationship between environmental variability and species diversity. For example, the Intermediate Disturbance Hypothesis (IDH) posits that environmental disturbances to competitive and successional processes will maximize species richness by preventing competitive exclusion (Chesson, 1994; Connell, 1978). In the case of apparent competition, analogized by immune suppression of two infecting parasites (Holt & Bonsall, 2017; Holt & Lawton, 1994), theory indicates that intermediate levels of disturbance will promote coexistence when a consumer has a larger impact on the stronger of two competitors (Wootton, 1998). More recent work suggests that within-host parasite diversity may be maintained by stabilizing and equalizing mechanisms (Bashey, 2015), including niche differentiation (Ramesh & Hall, 2023), competition-colonization trade-offs (T. A. Dallas et al., 2019), and environmental heterogeneity (Greenrod et al., 2024). Song and Spaak (2024) determined that in lower trophic levels, diversity is primarily constrained by fitness differences, which can be minimized via equalizing mechanisms like environmental variation (Song & Spaak, 2024; Spaak et al., 2021).

We considered the relationship between environmental variation and within-host parasite diversity along two axes: magnitude and frequency of stochastic disturbance (Mackey & Currie, 2001; Miller et al., 2011, 2012). First, we considered rising temperature variability as a feature of climate change (Lawson et al., 2015; Stocker et al., 2013; Vasseur et al., 2014). In ectothermic hosts, increasingly erratic external conditions will affect the internal conditions of the host, thereby also affecting any parasites the host may harbour (Cizauskas et al., 2017). A relationship between thermal variability and within-host parasite diversity may be of particular interest when considered alongside the long-debated relationship between climate change and disease-mediated amphibian declines (Daszak et al., 1999; Johnson, 2006; Lips et al., 2008; Rohr et al., 2008). While it has been posited that increasing temperatures may drive these declines, Rohr and Raffel (2010) demonstrated that El Niño events and associated regional temperature variability increase host susceptibility, and thus contribute to host population declines (Cohen et al., 2019; Rohr et al., 2008; Rohr & Raffel, 2010). Second, we considered environmental variability as a host trait. Bats (order *Chiroptera*) have long been acknowledged as prolific reservoirs for disease, and are unique among mammals in their ability to control body temperature in accordance with activity level (e.g. torpor, flight) (Brook & Dobson, 2015; Calisher et al., 2006; Irving et al., 2021; Luo et al., 2021). It has been theorized that thermal variation may contribute to the efficacy of bats’ immune systems, allowing them to carry microparasites without suffering increased morbidity (Fumagalli et al., 2021; Luo et al., 2021). Similarly, thermal variability may affect parasite coexistence. In combination, the effects of thermal variability on immune responses and parasite coexistence may help explain bats’ propensity to support large communities of parasites.

We used a model to investigate the impacts of environmental variability on the viability and maintenance of nonequilibrium parasite co-occurrence within hosts. We began by adapting Antia et al.’s (1994) model of within-host population dynamics to include two parasites that indirectly compete via the host’s immune system, and thus exhibit apparent competition (Holt, 1977; Råberg et al., 2006; Williamson, 1957). We analyzed deterministic, demographically stochastic, environmentally stochastic, and demographically- and-environmentally stochastic implementations of the model. Within the environmentally stochastic implementations of the model, we draw on the Metabolic Theory of Ecology to model the parasites’ replication rates as temperature-dependent, and vary the size and frequency of thermal disturbance within the system. Between and among the deterministic and stochastic implementations of the model, we compared the ability of the secondary parasite to invade a system containing the primary parasite, and the duration of time for which the primary and secondary parasites co-occurred. Further, we compared the total transmission of co-occurring parasites with the total transmission of a single parasite. We found that compared to a constant environment, a variable environment tended to increase the ability of the secondary parasite to invade the system, facilitated longer periods of co-occurrence between the primary and secondary parasites, and increased the transmissibility of co-occurring parasites. These results (1) suggest that climatic changes may have significant effects on parasite diversity within ectothermic hosts, (2) provide insight on a host’s ability to support large communities of parasites, and propose that variability within-hosts may influence transmission dynamics among-hosts.

## Methods

### Model Description

#### Within-Host Population Dynamics Model

We began with a model of within host population dynamics based on Antia et al., (1994) that represents the interaction between an infecting parasite and its host’s immune system as analogous to the interaction between a prey item and its predator. Per Antia et al. (1994), the density of the parasite population, *P*, is determined by its replication rate, *r*, and the rate at which it is cleared by the host’s immune system, *k*. The model assumes that the immune system’s replication rate is proportional to parasite density at low parasite densities, and saturates at high parasite densities, allowing the host’s immune system to either extirpate or succumb to the infection (Antia et al., 1994). We adapted the model to include a second parasite, *P*_*j*_, that indirectly competes with the primary parasite, *P*_*i*_, through the host’s immune response (Eqns. 1-3, Table 1). As such, the model describes an immune-mediated apparent competition (Holt, 1977; Råberg et al., 2006; Williamson, 1957). Parameters values were not chosen to describe a specific system, but to produce the dynamical behaviour described above and shown in Figure 1. Given that any number of parameter value combinations may produce this behaviour, we chose to maintain the initial conditions and select parameter values proposed by Antia et al. (1994). However, we increased the rate at which parasites are destroyed by the host’s immune system (*k*) and decreased the maximum replication rates of the primary and secondary parasites (*r*_*i*_, *r*_*j*_) to favour extirpation, and eliminate host death as an infection outcome (Table 1). We also conducted a sensitivity analysis to determine how variations in (1) the value of *k* and (2) the shape of the relationship between parasite replication rate and the temperature of the system would affect our results, and found the model to be robust (see pages 2, 4-8 of supplement for complete results).

**Table 1:** Parameter definitions and values.

| Symbol | Parameter | Values Stated in Antia et al. (1994) | Values Used |
| --- | --- | --- | --- |
| $P_i$ and $P_j$ | Parasite abundance | 1* | 1* |
| $I$ | Host immune cell abundance | 1* | 1* |
| $r_i$ and $r_j$ | Replication rates of parasites | 0.1 – 10.0 | 0.1 – 2.0<sup>†</sup> |
| $k$ | Rate at which parasites are destroyed by host's immune system | $10^{-3}$ | $10^{-2}$ |
| $\rho$ | Maximum replication rate of host's immune system | 1.0 | 1.0 |
| $\phi$ | Parasite density at which replication rate of host's immune system is half its maximum | $10^3$ | $10^3$ |
| $D$ | Lethal within-host parasite density | $10^9$ | $10^9$ |

\* Initial conditions
<sup>†</sup> Dependent on the temperature of system in environmentally stochastic simulations

**Figure 1:**
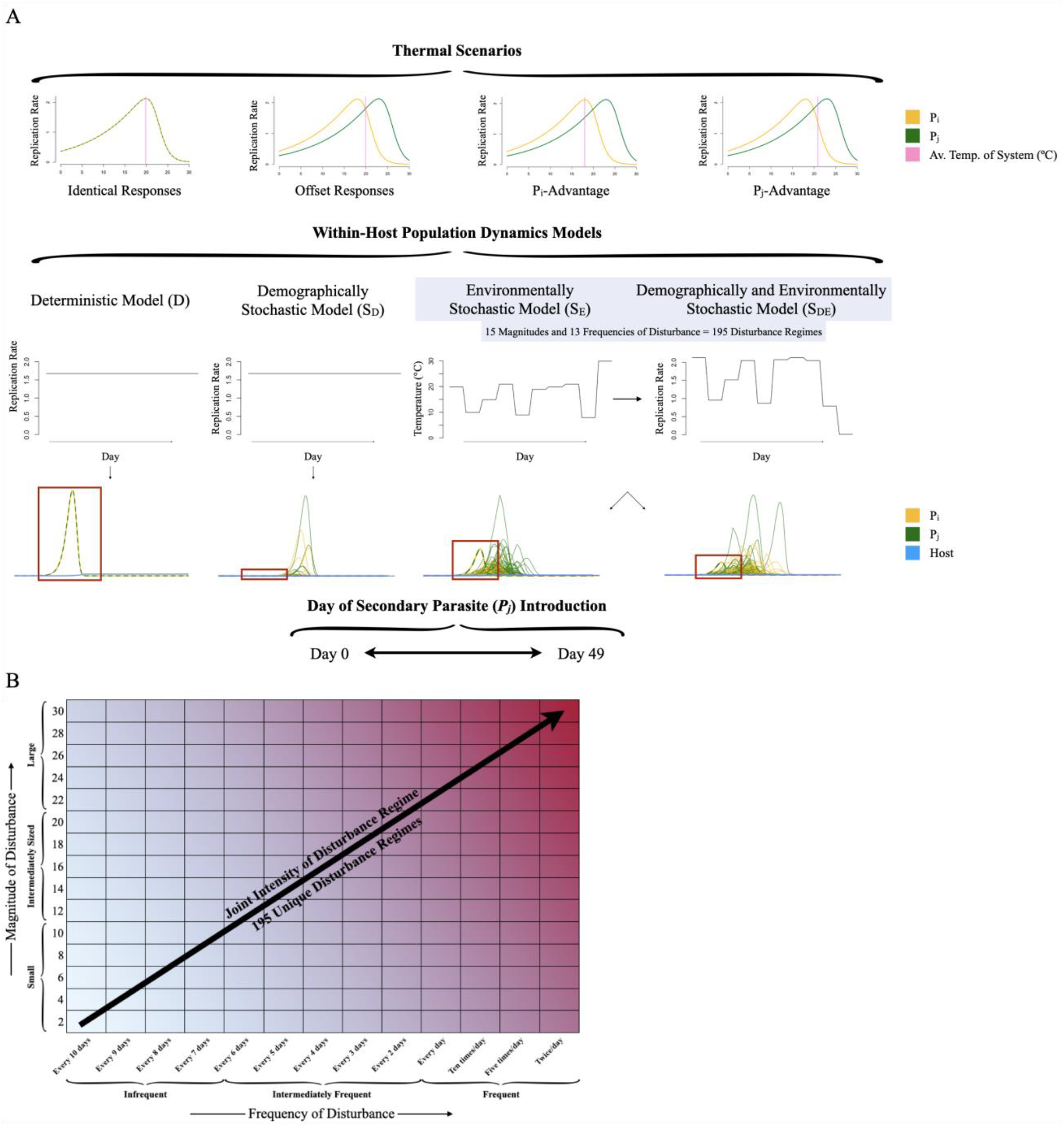
(A) Shows a schematic detailing simulation experiment. When simulating each of four models (deterministic, demographically stochastic, environmentally stochastic, and demographically and environmentally stochastic), we considered four thermal scenarios (identical responses, offset responses, *P*_*i*_-advantage, and *P*_*j*_-advantage). When simulating the environmentally stochastic models (in blue), we considered 15 sizes and 13 frequencies of disturbance, totalling 195 unique disturbance regimes. We additionally considered 50 days on which the secondary parasite could be introduced. We conducted 1000 simulations per stochastic model and treatment. Examples of such simulations are shown as time series. (B) Shows a schematic detailing how size and frequency interact to produce disturbance regimes.

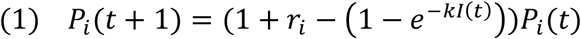

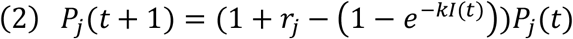

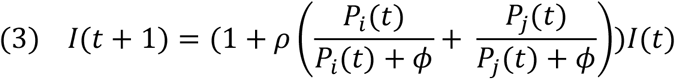

To compare the impacts of different types of stochasticity on parasite diversity, we wrote deterministic (D), demographically stochastic (S_D_), environmentally stochastic (S_E_), and demographically- and-environmentally stochastic (S_DE_) implementations of the model. Within the demographically stochastic implementation of the model, we wrote the model’s replication terms (*r*_*i*_·*P*_*i*_(*t*), *r*_*j*_·*P*_*j*_(*t*), and 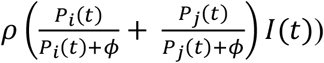 as Poisson processes, and mortality term (1 − *e*^−*kI*(*t*)^) as a binomial process

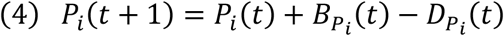

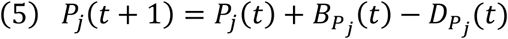

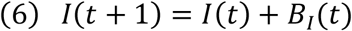

In Eqns. (4) and (5), 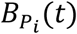 and 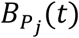 are Poisson random variables such that 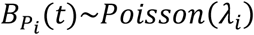, where *λ*_*i*_=*r*_*i*_·*P*_*i*_(*t*), and 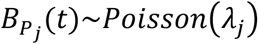, where *λ*_*j*_=*r*_*j*_·*P*_*j*_(*t*). In Eqn. (6), *BI*(*t*) is a Poisson random variable such that *I* (*t*)∼*Poisson*(*λ*), where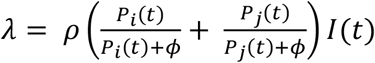. In Eqns. (4) and (5), 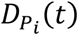 and 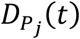 are binomial random variables such that 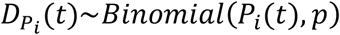, where *p*=1 − *e*^−*kI*(*t*)^, and 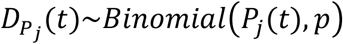. To write the environmentally stochastic implementation of the model, we used randomly generated temperature series to dictate the temperature of the system (*T*), and allowed the replication rates of each parasite (*r*_*i*_, *r*_*j*_) to vary in response (Eqns. 7-9). We chose to write *r*_*i*_ and *r*_*j*_, though not *k*, as temperature-dependent to isolate the effect of thermal variability on traits that can be metabolically scaled, and constrain the scope of the work. We did not write *ρ*or *ϕ*as temperature-dependent to constrain the scope of the work.

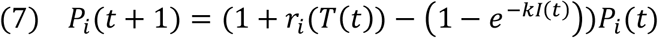

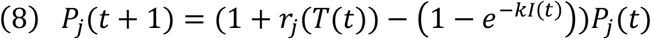

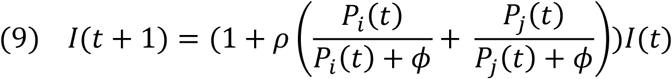

We used first-order autocorrelated Gaussian noise to generate temperature series. We initiated the series with a value of zero, and generated the next value by multiplying the previous value by the coefficient of autocorrelation (0.25), and summing the resulting value with a value drawn from a normal random variable with a mean of zero. We varied the standard deviation of the normal random variable to control the size of disturbance, as described below. We then added the average temperature of the system to each value in the series. Resulting values outside the thermal range of the system were re-drawn. We then converted the temperature series into replication rates using various parameterizations of the Sharpe-Schoolfield sub-model, as described below (Eqns. S1.1, S1.2; Table S2).

#### Sharpe-Schoolfield Sub-Model

The Metabolic Theory of Ecology (MTE) predicts how metabolic rates control ecological processes across levels of organization, and explains how metabolic rates vary with body size and temperature (Brown et al., 2004). Building off the MTE, the Boltzmann-Arrhenius (BA) relation describes metabolism as a function of temperature. The Sharpe-Schoolfield model characterizes the performance of a trait as a function of temperature through a thermal performance curve (TPC) (Schoolfield et al., 1981). When applied to disease systems, the Sharpe-Schoolfield model has been shown to describe relationships between temperature and species traits, and has proven useful in investigating the relationship between rising temperatures and disease transmission.

We used various parameterizations of the Sharpe-Schoolfield model to generate paired thermal performance curves relating the temperature of the system to the replication rates of two parasites (*r*_*i*_ and *r*_*j*_) (Fig. S9). We generated four paired thermal performance curves, representing four thermal scenarios: (1) the identical responses scenario, (2) the offset responses scenario, (3) the *P*_*i*_-advantage scenario, and (4) the *P*_*j*_-advantage scenario (Fig. 1, Table S2). The identical responses scenario describes *r*_*i*_ and *r*_*j*_ as responding identically to the temperature of the system, and exhibits an average temperature of 19.9ºC (Fig. 1, Table S2, Fig. S9). The remaining scenarios describe a system within which *P*_*i*_ exhibits a higher replication rate at lower temperatures and *P*_*j*_ exhibits a higher replication rate at higher temperatures, and are differentiated by the average temperature of the system. The offset responses scenario exhibits an average temperature of 19.9ºC. At this temperature, the replication rate of *P*_*i*_ is equal to that of *P*_*j*_ (Fig. 1, Table S2, Fig. S9). The *P*_*i*_-advantage scenario exhibits an average temperature of 18ºC, giving *P*_*i*_ a thermal advantage over *P*_*j*_ (Fig. 1, Table S2, Fig. S9). We considered two versions, (I) and (II), of the *P*_*j*_-advantage scenario, each of which give *P*_*j*_ a thermal advantage over *P*_*i*_. (I) exhibits an average temperature of 20.9ºC, such that *P*_*j*_’s thermal advantage is comparable in size to that of *P*_*i*_, as described above (Fig. 1, Table S2, Fig. S9). (II) replicates the *P*_*i*_-advantage scenario, but re-assigns the low-temperature curve to *P*_*j*_ and the high-temperature curve to *P*_*i*_, to isolate the effects of asymmetry (Fig. S9). Given that (I) and (II) produced qualitatively comparable results, we will only discuss (I) in the main text (see supplement for complete results).

Given that the Sharpe-Schoolfield model cannot generate negative values, it cannot be used to explore cases in which extreme temperatures result in parasite mortality and population decline. To compensate, we developed an alternative set of paired thermal performance curves, through which extreme temperatures produce “negative” replication rates, and spur population decline (see “Extended analysis” in supplement; Fig. S19).

### Model Simulations

Within each thermal scenario, we simulated each model (D, S_D_, S_E_, S_DE_) for fifty days. Simulating for fifty days proved sufficient to capture the full course of parasite infection and clearance across all treatments. We began each set of simulations with the primary parasite and the host’s immune system in the system, at initial abundances of one. We then considered all cases for the introduction of the secondary parasite from its introduction at the onset of infection, to its introduction on the forty-ninth day of infection.

For each environmentally stochastic model (S_E_, S_DE_) simulation, we generated a new temperature series and two corresponding series of replication rates. When determining how new temperatures would be selected, we considered two axes of thermal disturbance: size and frequency. We controlled the size of disturbance by changing the standard deviation of the normal random variable in the temperature selection function, starting from two and increasing by two until the effect was fully characterized at a value of thirty, resulting in fifteen sizes of disturbance (Fig. 1B). We considered standard deviations between two and ten to denote small disturbances, between twelve and twenty to denote intermediately sized disturbances, and between twenty-two and thirty to denote large disturbances (Fig. 1B). To control the frequency of disturbance, we re-assigned the temperature of the system every ten, nine, eight, seven, six, five, four, three, and two days, once per day, and two, five, and ten times per day to fully characterise the effect, resulting in thirteen frequencies of disturbance (Fig. 1B). We considered re-assignments every ten to seven days to denote infrequent disturbances, every six to two days to denote intermediately frequent disturbances, and every day to ten times per day to denote frequent disturbances (Fig. 1B). Considering size and frequency together resulted in 195 unique disturbance regimes. We conducted 1000 simulations per stochastic model and treatment (Fig. 1A). Two hundred such simulations are shown in Figures 1A and S10.

### Co-Occurrence Analyses

Given successful invasion by the secondary parasite (*P*_*j*_(*t* + 1) > *P*_*j*_(*t*)), we defined longevity of co-occurrence as the amount of time for which both parasites exhibited population abundances equal to or greater than one.

We summarised these results as distributions for each stochastic model and treatment. Using the S_D_ model as a baseline, we investigated the effects of environmental variability on the viability and maintenance of parasite diversity by comparing the ability of the secondary parasite to invade a single-parasite system, and the duration of time for which the primary and secondary parasites co-occurred across stochastic treatments. Additionally, we analysed (1) the proportion of stochastic simulations per model and treatment that resulted in longer periods of co-occurrence relative to the deterministic model, and (2) the competitive outcomes of each treatment to gain insight on underlying mechanisms.

### Transmissibility Analyses

To explore the impacts of co-infection on transmission potential, we compared the lifetime replicative outputs of co-occurring parasites with those of a single parasite in constant and variable environments. As per Antia et al., (1994), we determined the total transmission (*q*_*i*_, *q*_*j*_, *q*, where *q* is the total transmission of a single parasite) of a parasite (*P*_*i*_, *P*_*j*_, *P*, where *P* is a single parasite) from an infected host by taking the integral of the rate of transmission (*u*) of each parasite from the host over the duration of infection (Eqns. 10.1-10.3). For simplicity, we set the rate of transmission to one.

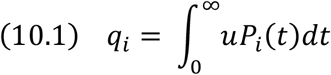

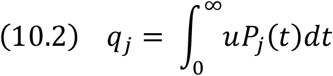

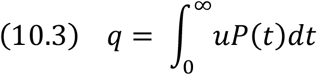

To compare constant and variable environments, we analysed multi-parasite and single-parasite versions of the S_D_ and S_DE_ models.

## Results

Analytically, the secondary parasite could invade (*P*_j_ (*t* + 1) > *P*_j_ (*t*)) when 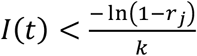. Our parameterization of the S_D_ model allowed the secondary parasite to invade when introduced before the tenth to twelfth day of infection, resulting in 8.4 to 9.2 days of co-occurrence, depending on thermal scenario (Fig. S11, Supplementary Datafile 1-1:10).

### Co-occurrence Analysis

Simulation outputs show that compared to a deterministic baseline, the addition of demographic stochasticity has little effect on co-occurrence (Supplementary Datafile 2-1:10), while the addition of thermal variability increases the ability of the secondary parasite to invade the system and facilitates longer periods of co-occurrence between parasites. While longevities of co-occurrence are maximised under the *P*_*j*_-advantage scenario, the difference between deterministic and environmentally stochastic implementations is not consistently maximised by any one thermal scenario. Additionally, while larger and more frequent disturbances tend to promote longer periods of co-occurrence, increasingly large disturbances produce a less steep effect than do increasingly frequent disturbances. These findings are common among thermal scenarios.

When both parasites were introduced simultaneously, S_D_ model simulations resulted in periods of co-occurrence that were, on average, 4.5% to 3.1% shorter (0.4 to 0.3 days) than deterministically simulated periods of co-occurrence (Figs. 2, S12). In comparison, simulations of the environmentally stochastic models (S_E_, S_DE_) resulted in periods of co-occurrence that were up to 46.2% (S_E_) and 36.0% (S_DE_) longer (3.9 (S_E_) and 3.0 (S_DE_) days) than deterministically simulated periods of co-occurrence (Figs. 2, S12). Across all 195 disturbance regimes, simulations of the S_E_ model resulted in mean periods of co-occurrence that were between 2.7% shorter (0.2 days) and 46.2% longer (3.9 days) than their deterministic counterparts (Figs. 2, S12). The identical responses scenario promoted the shortest and longest mean periods of co-occurrence (Figs. 2-3, S12-S13). Simulations of the S_DE_ model resulted in mean periods of co-occurrence that were between 6.8% shorter (0.6 days) and 36.0% longer (3.0 days) than deterministically simulated periods of co-occurrence (Figs. 2, S12). Again, the identical responses scenario promoted the shortest and longest mean periods of co-occurrence (Figs. 2-3, S12-S13). Across thermal scenarios, mean periods of co-occurrence were shortest when disturbances were large (standard deviation of the normal random variable in the temperature autocorrelation function = 22 to 30) and infrequent (once every 7 to 9 days) (Figs. 3, S13). Mean periods of co-occurrence were longest when disturbances were large (SD = 24 to 30) and frequent (five to ten times per day) (Figs. 3, S13).

**Figure 2:**
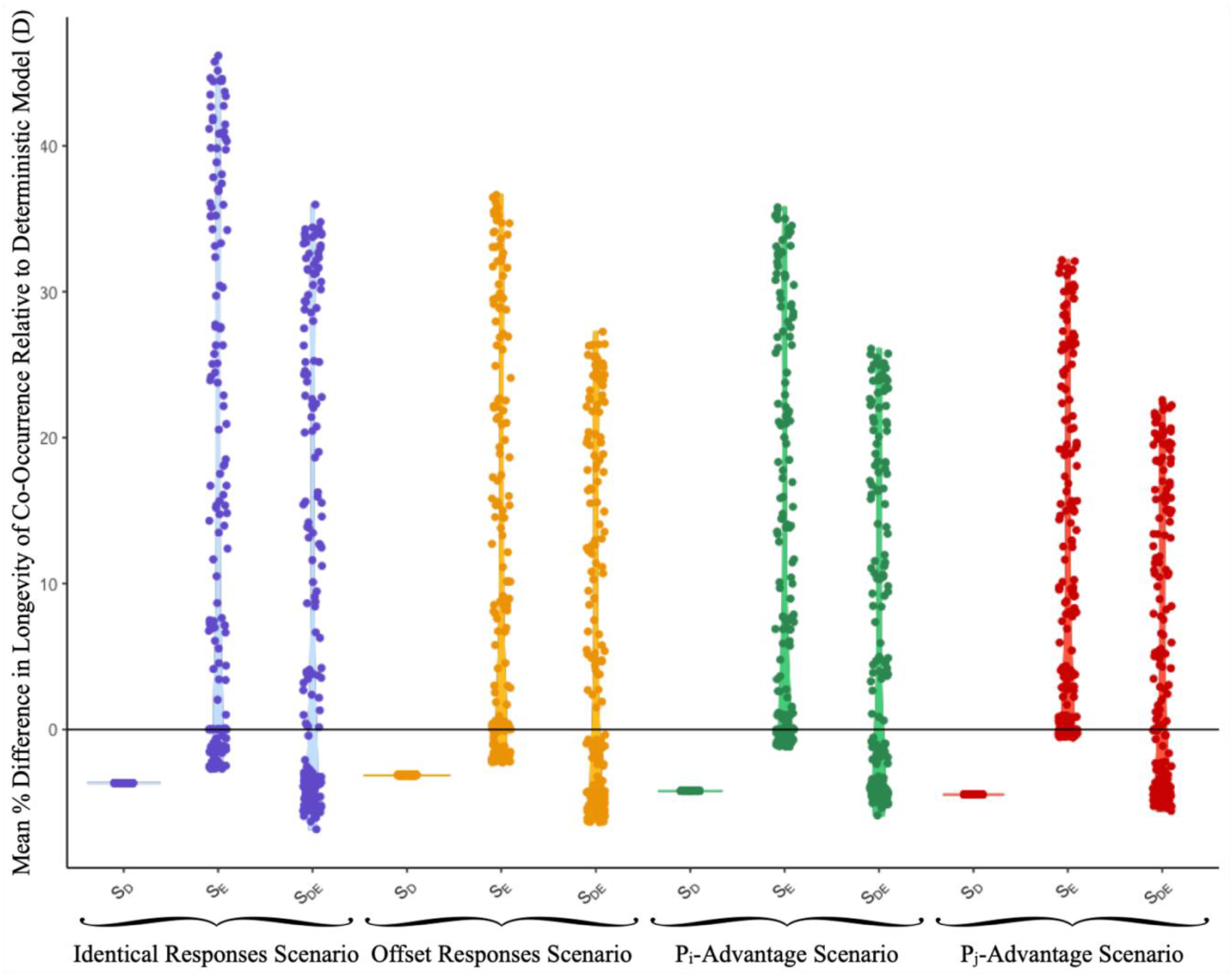
The mean percent difference in longevity of co-occurrence relative to deterministically simulated periods of co-occurrence, when both parasites are introduced at the same time. Single points above “S_D_” labels represent the mean percent difference in longevity of co-occurrence relative to deterministic thresholds across 1000 demographically stochastic simulations. Each point above “S_E_”/”S_DE_” labels represents the mean percent difference in longevity of co-occurrence relative to deterministic thresholds across 1000 environmentally stochastic simulations. As such, there are 195 points above each “S_E_”/”S_DE_” label, corresponding to 195 environmental disturbance regimes.

**Figure 3:**
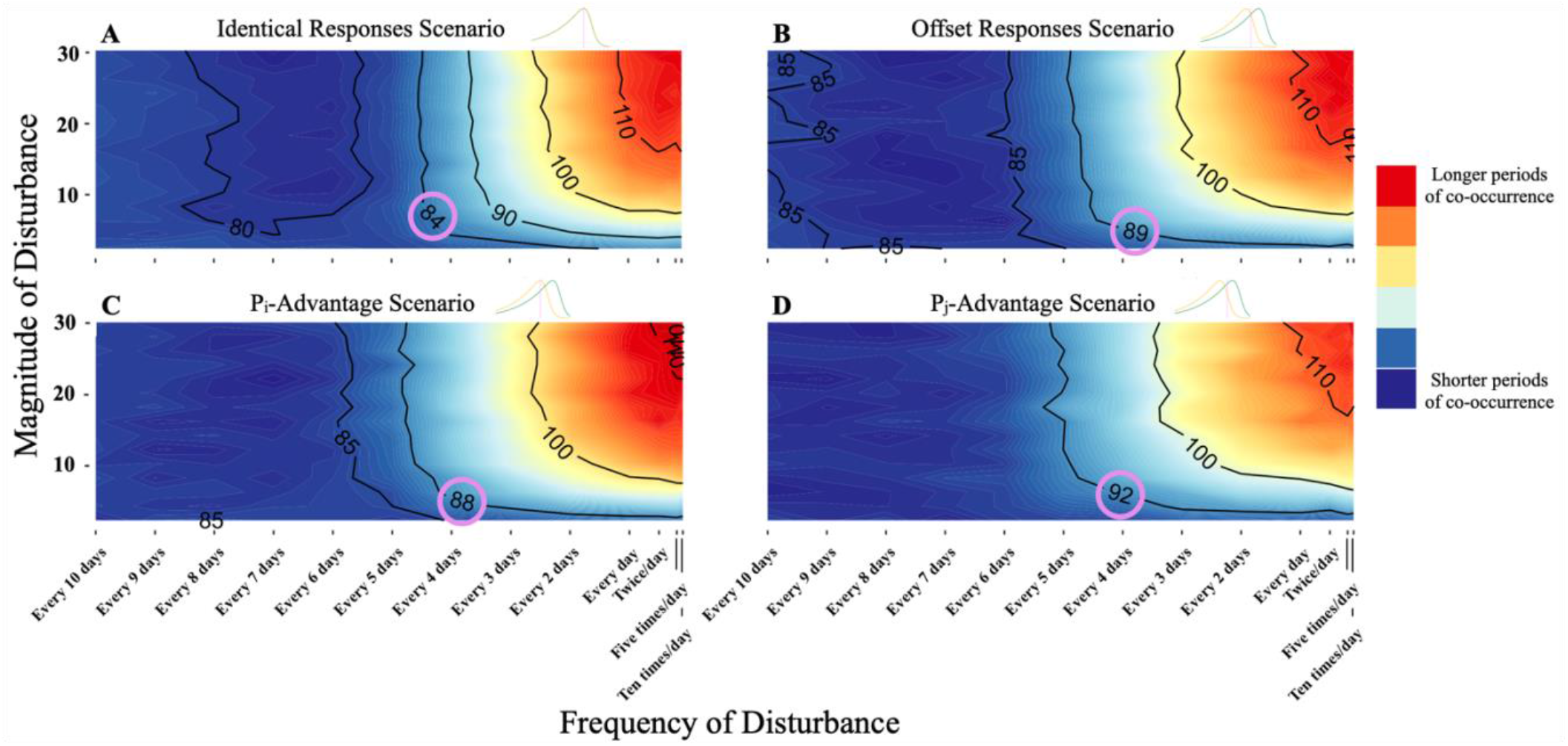
Fully factorial representations of the impacts of varyingly sized and timed disturbances on the longevity of co-occurrence. The plot shows the mean number of time steps (10 time steps/day) for which *P*_*i*_ and *P*_*j*_ have population abundances greater than one as per the S_DE_ model. The circled numbers denote the number of time steps for which *P*_*i*_ and *P*_*j*_ can co-occur, per the deterministic model.

More generally, as disturbances increased in size, periods of co-occurrence increased in length towards a saturation point (standard deviation of the normal random variable in the temperature autocorrelation function ≈ 24-30, depending on thermal scenario) (Figs. S14, S18, Supplementary Datafiles 3-1:5, 4-1:5). However, an analysis of simulations within which an alternative set of TPCs allowed for extreme temperatures to cause population decline revealed a parabolic relationship between disturbance size and longevity of co-occurrence, that peaked when disturbances were intermediately sized (standard deviation of the normal random variable in the temperature autocorrelation function ≈ 8-16) (Fig. S20).

When disturbances were intermediately frequent (once every two to six days), periods of co-occurrence were similar in length to deterministically simulated periods of co-occurrence. Interestingly, when disturbances were infrequent (once every six to ten days), mean periods of co-occurrence were shorter than deterministically simulated periods of co-occurrence (Fig. S14, S21). Per the deterministic model, *P*_*i*_ and *P*_*j*_ can only co-occur for 8.4 to 9.2 days. Thus, a system subjected to infrequent disturbances is functionally similar to the deterministic system, making the outcome of infection reliant on initial conditions. When disturbances were frequent (ten times per day to once per day), mean periods of co-occurrence were longer than deterministically simulated periods of co-occurrence. Within a given disturbance regime, longevities of co-occurrence were normally distributed (Figs. S14-S15, S23).

### Rarity and Competitive Outcomes

When the secondary parasite was introduced before or on the invasion threshold (10^th^ or 12^th^ day of infection, depending on thermal scenario), 29%-71% (S_E_) and 46%-58% (S_DE_) of environmentally stochastic simulations resulted in longer periods of co-occurrence relative to deterministic simulations (Fig. 4). When the secondary parasite was introduced after the invasion threshold, the likelihood that an environmentally stochastic simulation resulted in some period of co-occurrence sharply declined. When the secondary parasite was introduced one to twelve days after the invasion threshold, 1%-31% (S_E_) and 1%-46% (S_DE_) of environmentally stochastic simulations resulted in some period of co-occurrence (Fig. 4). Thus, environmental variability extended the invasion threshold by 50%-75% (S_E_) and 100%-120% (S_DE_), depending on thermal scenario (Fig. 4).

**Figure 4:**
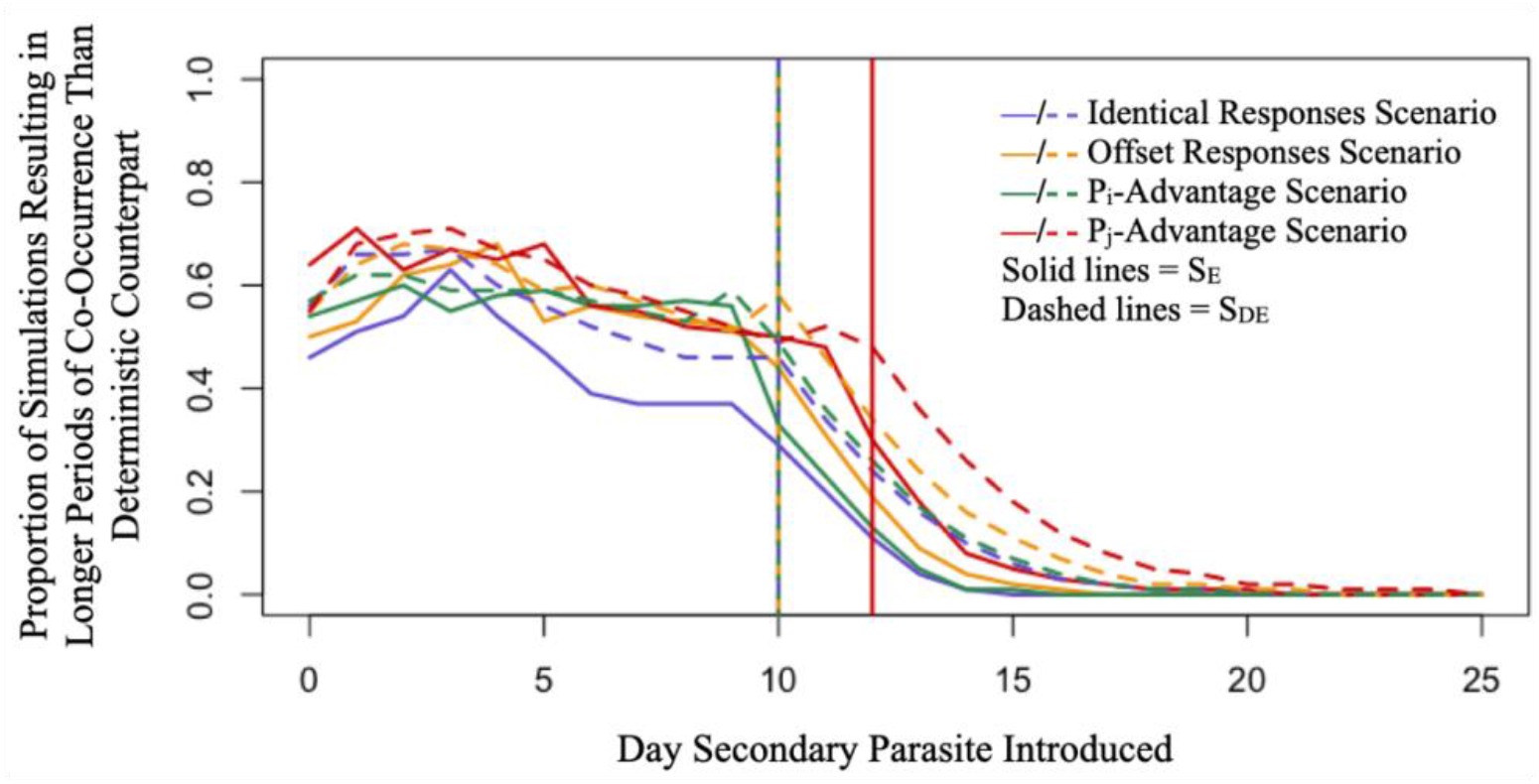
Proportion of environmentally stochastic simulations (SD = 10, frequency of disturbance = once per day) resulting in longer periods of co-occurrence relative to deterministic simulations by the day on which the secondary parasite was introduced. Vertical lines show the first day on which the introduction of the secondary parasite does not result in any co-occurrence between the two parasites, as per the deterministic model (D).

Analyzing competitive outcomes at intermediate levels of disturbance (SD = 10, frequency of disturbance = once per day) revealed that under the offset responses scenario, when *P*_*i*_ and *P*_*j*_ were introduced at the same time, *P*_*j*_ outlasted *P*_*i*_ 65.5% (S_E_) and 63.4% (S_DE_) of the time (Fig. S22-S23). Additionally, under the *P*_*i*_-advantage scenario, when *P*_*i*_ and *P*_*j*_ were introduced at the same time, *P*_*i*_ and *P*_*j*_ were equally likely to outlast one another (*P*_*j*_ outlasts *P*_*i*_ 49.9% (S_E_) and 51.0% (S_DE_) of the time), but under the *P*_*j*_-advantage, *P*_*j*_ was much more likely to outlast *P*_*i*_ (*P*_*j*_ outlasts *P*_*i*_ 76.6% (S_E_) and 70.4% (S_DE_) of the time) (Fig. S21-S23). Across thermal scenarios, when the secondary parasite, *P*_*j*_, was introduced in the middle of the infection period, *P*_*i*_ was more likely to outlast *P*_*j*_. When *P*_*j*_ was introduced late in the infection period, *P*_*j*_ was more likely to outlast *P*_*i*_ (Fig. S22-S23). Regardless of when *P*_*j*_ was introduced, simulations resulting in especially long periods of co-occurrence are equally likely to show either parasite outlasting the other (Fig. S22-S23).

### Transmissibility

Across thermal scenarios, simulations of the multi-parasite S_D_ model produced lifetime replicative outputs (LROs) that were one to two orders of magnitude lower than those produced by simulations of the single-parasite S_D_ model (Supplementary Datafiles 5-1:16). Similarly, under the first three thermal scenarios, simulations of the multi-parasite S_DE_ model produced LROs that were one to two orders of magnitude lower than those produced by simulations of the single-parasite S_DE_ model. However, under the *P*_*j*_-advantage scenario and among 62 of 195 (32%) disturbance regimes, the average transmission potential of the co-occurring secondary parasite, *P*_*j*_, exceeded that of a single parasite (Fig. 5). Among these disturbance regimes, the transmission potential of *P*_*j*_ exceeded that of a single parasite by an average of 2% to 135% (Supplementary Datafiles 5-8). The transmission potentials of co-occurring parasites peaked when disturbances were infrequent (once every ten days) to intermediately frequent (once every six days) (Fig. S24). While intermediately large and frequent disturbances maximally increased longevities of co-occurrence, such infections did not result in particularly high transmission potentials, indicating that especially long periods of co-occurrence may be facilitated by lower lifetime replicative outputs that do not trigger a large immune response, and thus allow for more persistent, but less transmissible, infections.

**Figure 5:**
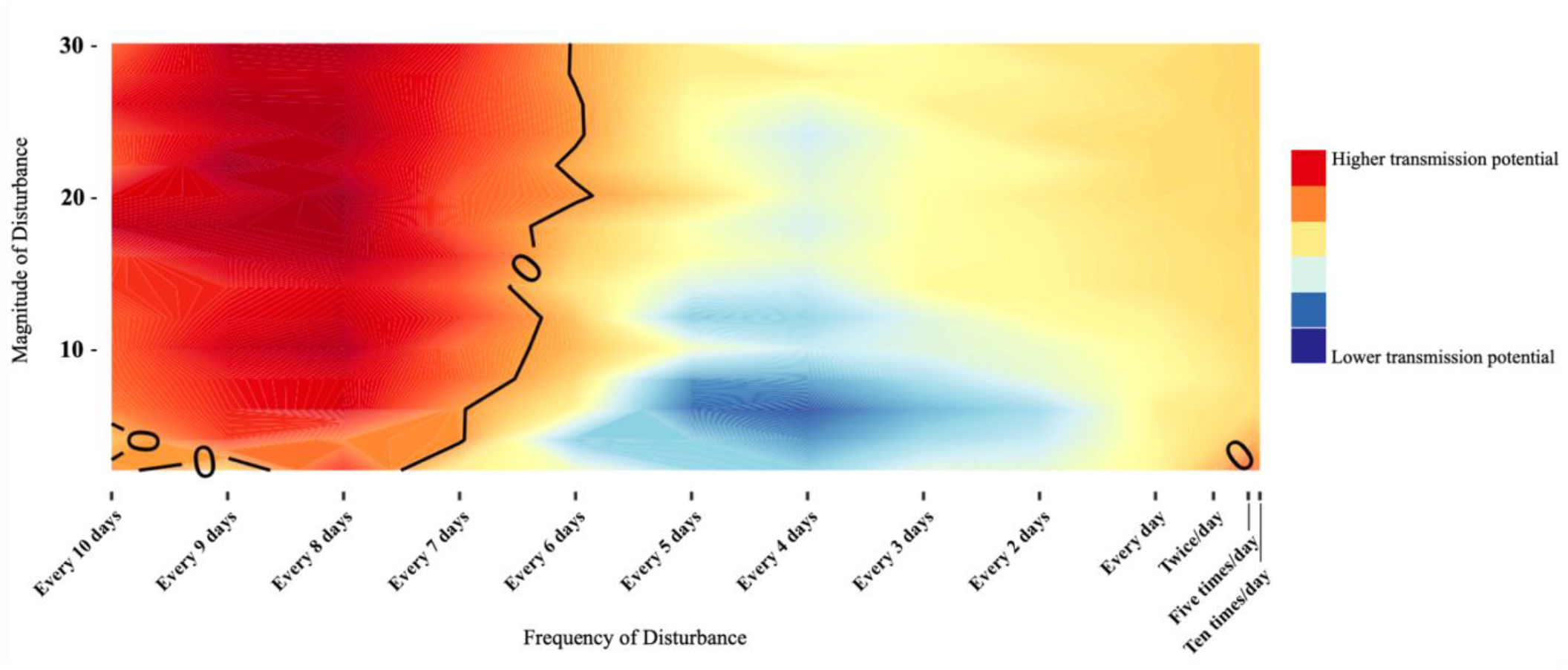
The impacts of environmental variability on transmission potential under the *P*_*j*_-advantage scenario. The plot shows a fully factorial representation of the impacts of thermal disturbance on the transmission potential of *P*_*j*_, relative to the equivalent single-parasite treatment. The black line indicates disturbance regimes under which the average transmission potential of *P*_*j*_ was equal to that of a single parasite. Colours warmer than denoted by the black line indicate disturbance regimes under which the average transmission potential of *P*_*j*_ exceeded that of a single parasite; colours cooler than denoted by the black line indicate the opposite.

## Discussion

We investigated the potential for environmental variability to affect the viability and maintenance of within-host parasite diversity. Our work provides theoretical evidence that relative to a constant environment, a variable environment can increase the ability of a secondary parasite to invade a single-parasite system, promote longer periods of co-occurrence between parasites, and increase the transmissibility of co-occurring parasites. In more detail, thermal disturbances of various sizes and frequencies allowed a secondary parasite to invade a single-parasite system up to twelve days after the deterministic invasion threshold, extending the threshold by 100% to 120% (S_DE_ model, Fig. 4). Large and frequent thermal disturbances enabled parasites to co-occur for up to 37.8% longer, relative to a constant environment (S_DE_ model, Figs. 2, S12). Finally, under the *P*_*j*_-advantage scenario and among 32% disturbance regimes, the average transmission potential of the co-occurring secondary parasite exceeded that of a single parasite (S_DE_ model, Fig. 5).

Across thermal scenarios, the relationship between disturbance size and longevity of co-occurrence was initially positive and eventually saturating (Fig. S14, S18). Large disturbances produced a saturating response rather than the expected parabolic response due to the structure of the Sharpe-Schoolfield sub-model. Allowing for extreme temperatures to result in parasite mortality and population decline produced a parabolic response, implying that intermediate sizes of disturbance maximised the viability and longevity of parasite co-occurrence (Figs. S19-S20). When thermal disturbances were small, the parasite with the higher initial replication rate was more likely to grow quickly and trigger a commensurate immune response that would competitively exclude the other. As thermal disturbances increased in size, it became increasingly likely that when the temperature of the system changed, the competitive advantage would shift from one parasite to the other, minimizing fitness differences over the course of an infection and promoting co-occurrence. However, per the extended analysis, large thermal disturbances caused parasite populations to alternatively explode and crash, triggering a sizable immune response, increasing the likelihood of extirpation, and precluding co-occurrence.

The relationship between climate change and disease-mediated amphibian declines has long been debated. While some argue that climatic changes may impede immune function or result in performance gaps between parasites and their hosts, making amphibians more susceptible to infection, others argue that insufficient evidence should preclude a causal link (Cohen et al., 2019; Rohr et al., 2008). While much of the literature proposes mean-climate hypotheses, some evidence suggests that intensifying El Niño events are increasing thermal variability, which may in turn affect amphibians’ susceptibility to or ability to defend against infection (Rohr & Raffel, 2010). Congruently, our findings provide theoretical evidence that thermal variability promotes the viability and maintenance of within-host parasite diversity, and thus suggest that as the climate continues to change, attention should be paid to shifts in microparasite communities and their impacts on ectothermic hosts. In particular, well-documented climate-driven increases in disease incidence among a wide variety of marine organisms may constitute an important starting point for future work (Bates et al., 2009; Bruno et al., 2007; Tracy et al., 2019).

Across thermal scenarios, low and high frequencies of disturbance impeded and promoted the viability and longevity of parasite co-occurrence, respectively (Figs. 3, S13-S14, S21). Given the transient nature of infection specified by the underlying model, infrequent disturbances (once every seven to ten days) resulted in the temperature of the system being disturbed towards the end of the infection period, when parasite populations were small and especially susceptible to unfavourable conditions. Conversely, frequent disturbances (ten times per day to once per day) resulted in the temperature of the system being disturbed throughout the duration of the infection period, and as such, neither favourable nor unfavorable conditions lasted long enough for one parasite to gain a decisive advantage over and exclude the other, and thus promoted extended periods of co-occurrence. While it has been theorized that internal temperature variation may aid bats’ immune responses, allowing them to carry microparasites without suffering outsized morbidity (Fumagalli et al., 2021), these findings suggest that internal temperature variation may also facilitate parasite co-occurrence within bats, and thus may help explain bats’ propensity to support diverse communities of parasites.

Under the offset responses scenario, *P*_*j*_ often outlasted *P*_*i*_, while under the *P*_*i*_-advantage scenario, *P*_*i*_ and *P*_*j*_ were equally likely to outlast each other (Fig. S22-S23). Under the *P*_*j*_-advantage scenario, *P*_*j*_ was much more likely to outlast *P*_*i*_ (Fig. S22-S23). These outcomes are consequences of the asymmetrical structure of the thermal performance curves, which made unfavourable conditions more disadvantageous for *P*_*i*_ than *P*_*j*_ (Sinclair et al., 2016). Further, regardless of when *P*_*j*_ was introduced, simulations resulting in especially long periods of co-occurrence were equally likely to show either parasite outlasting the other. Taken together, these results suggest that to predict the competitive outcome of a multiple infection, one may need to consider the interplay between how well adapted a parasite is to its host, priority effects, and the responses of biological rates to environmental factors. While one might expect a specialist or well-adapted parasite to exploit its host more efficiently than a generalist or novel parasite, the results of this work suggest that environmental variability can even the scales, and allow a generalist or novel parasite to establish and persist, as predicted by the intermediate disturbance hypothesis (Chesson, 1994; Connell, 1978; Roxburgh et al., 2004). Similarly, while one might additionally expect a colonizing parasite to benefit from an initial lack of competition, the results of this work again suggest that environmental variability and disturbance may contribute to the structuring of communities by allowing latecomers to establish and persist more often and for longer than they otherwise would (Jiang & Patel, 2008; Symons & Arnott, 2014).

While the metabolic theory of ecology posits that certain biological rates (e.g. metabolic rate, replication/growth rate, etc.) will exhibit an asymmetrical and unimodal response to temperature, other vital rates may exhibit symmetrical or bimodal responses to temperature (Dell et al., 2014; Molnár et al., 2017; Sinclair et al., 2016). This work suggests that the shape of a thermal performance curve can meaningfully affect the nature and outcome of an infection. The shape of a thermal performance curve can also dictate the effects of stochasticity on persistence. Here, thermal variation pushes the system away from a parasite’s optimal temperature, and thus, away from its maximum replication rate. In accordance with Jensen’s inequality, compared to a deterministic system, the introduction of stochasticity results in a lower average population growth rate (Denny, 2017; Jensen, 1906; Vasseur et al., 2014). In the absence of thermal dependency, the introduction of stochasticity will also result in a lower average population growth rate over time (Tuljapurkar, 1990). In this system, lower average parasite replication rates slow down the infection and subsequently, the immune response, allowing parasites to co-occur for longer. As anthropogenic changes alter host-parasite interaction networks, understanding how evolutionary and environmental factors combine to affect interactions between parasites will be essential in predicting and managing spillover and emergence in wildlife and human populations (Paull et al., 2012; Pedersen & Fenton, 2007).

Developing an understanding of how the impacts of environmental variability on parasite diversity within hosts may scale to affect among-host processes and events, including transmission, spillover, and emergence will be essential to the maintenance of wildlife, human, and ecosystem health (Allocati et al., 2016; Halpin et al., 2000; Johnson, 2006). While increased within-host parasite diversity provides more strains or species of a parasite or parasites an opportunity to transmit, within-host competition may result in decreased parasite loads, thus decreasing the transmission probability of any one strain or species, and altering expected transmission patterns (Johnson et al., 2013; Pedersen & Fenton, 2007). Similarly, our findings provide theoretical evidence that when environmental variations are large and frequent, within-host competition decreases the total transmission of co-occurring parasites. As such, while large and frequent disturbances may maximally increase the amount of time for which two parasites co-occur, such infections do not result in particularly high transmission potentials, indicating that especially long periods of co-occurrence may be facilitated by lower parasite replication rates that do not trigger a large immune response, and thus allow for more persistent infections. However, we found that approximately a third of tested disturbance regimes allowed the replicative outputs of co-occurring parasites to exceed those of single parasites, indicating that the effects of environmental variability may not be limited to the promotion of within-host parasite diversity, but may also promote transmission between hosts (Fig. 5, S24). Disturbances of intermediate size and low to intermediate frequency may constitute a sizable region within which environmental variation can decrease fitness differences between parasites and promote within-host parasite diversity, while allowing parasites to achieve sufficiently high parasite loads, resulting in more diverse and transmissible infections (Fig. 5, S24).

While both Antia et al.’s (1994) model and our modified model should be broadly applicable to an array of within-host multi-parasite systems, including multi-strain or cross-species, both models are relatively simple as to remain generalizable, and thus cannot represent chronic infections, systems within which parasites exhibit more complex interactions with their host or their host’s immune system, nor systems within which parasites directly interact with one another. As such, it may be interesting to validate the theoretical insights of this work in an empirical system. This work is also limited by the scope with which thermal dependency was considered. Given the thermal sensitivity of immune function in ectotherms and endotherms, it may be worthwhile to consider the thermal dependency of immune processes like response time and replication rate in tandem with the thermal dependency of parasite replication rates (Butler et al., 2013; Wright & Cooper, 1981). Similarly, using nonlinear averaging to assess the response of a nonlinear process to variability is contentious. While many theoretical works and some empirical studies support the use of nonlinear averaging and find evidence for Jensen’s inequality, borne out as disproportionate effects of thermal variability on performance, many empirical studies have contested an implicit assumption of nonlinear averaging: that traits should respond instantaneously to change, without acclimation (Bernhardt et al., 2018; Estay et al., 2011; Krichel et al., 2023; Sinclair et al., 2016). Thermal acclimation through phenotypic plasticity alters thermal performance curves, and as such, produces a more complex relationship between temperature and biological rates that may not be well-characterized by nonlinear averaging (Schulte et al., 2011; Sckrabulis et al., 2022). With regards to host-parasite interactions, the temperature variability hypothesis posits that owing to their small relative size, parasites should acclimate more quickly than their hosts, though the speed at which various parasites may acclimate remains unclear (Raffel et al., 2013; Rohr et al., 2013). Interestingly, some empirical studies of the impacts of environmental variability on host-parasite interactions have considered thermal acclimation, yielding a range of conclusions including decreased pathogen prevalence, reduced infection prevalence, and decreased resistance in hosts (T. Dallas & Drake, 2016; Duncan et al., 2011; Raffel et al., 2013).

Ecological theory has long predicted that environmental variability supports species diversity, from the IDH (Chesson, 1994; Connell, 1978; Huston, 2014; Roxburgh et al., 2004) to modern coexistence theory (Barabás et al., 2018; Chesson, 1994, 2000, 2003; Chesson & Huntly, 1997). Work on trophic interactions, wherein apparent competition is considered analogous to interactions between the host’s immune system and infecting parasites (Holt & Bonsall, 2017; Holt & Lawton, 1994), indicates that intermediate levels of disturbance can promote coexistence (Wootton 1998) and that within lower trophic levels (e.g. among competing parasites), diversity is primarily constrained by fitness differences, which can be minimized via equalizing mechanisms like environmental variation (Song & Spaak). Here, we provide theoretical evidence that the predicted relationship between environmental variability and species diversity is borne out in within-host disease systems. Camus wrote that plagues, despite their commonality, find us unprepared (Camus, Albert, 2008). Improving our understanding of how environmental variability affects parasite diversity within- and between-hosts will expand our understanding of the mechanisms that can facilitate disease spread and emergence, and allow us to make robust predictions, and develop and implement mitigation strategies as we move into an increasingly uncertain future. In brief, we will be found *less* unprepared.

## Acknowledgements

This work was supported by The Emerging and Pandemic Infections Consortium Doctoral Award (MJ-C), and Ontario Graduate Scholarship (MJ-C), a Natural Sciences and Engineering Research Council of Canada Discovery Grant (MK), and a Canada Research Chair (MK).

## Author Contributions

MJC and MK conceived the ideas and designed the methodology; MJC collected the data; MJC analysed the data; MJC led the writing of the manuscript. All authors contributed critically to drafts and gave final approval for publication.

## Data Availability Statement

Time series data available from the Federated Research Data Repository at https://doi.org/10.20383/103.0764. Scripts and replication materials available from GitHub at https://github.com/MadelineJC/EnvironmentalVariability_Repo.

