## Supplemental Information for "Environmental variability can promote parasite diversity within hosts and transmission among hosts"

**Submitted To: Theoretical Ecology**

[1]Madeline Jarvis-Cross (ORCID: 0009-0008-0527-9369)
[1]Martin Krkosek (ORCID: 0000-0001-7591 7954)

1 The University of Toronto, Department of Ecology and Evolutionary Biology
25 Willcocks Street, Toronto, ON, M5S 3B2

**Table of Contents**

Sensitivity analysis (Fig. S1, Table S1) 2
Extended analysis 3
High-average temperature environment analysis (Figs. S2-S8) 4
Sharpe-Schoolfield thermal sub-model 8
Table S2: Parameter definitions and values for Sharpe-Schoolfield sub-model 8
Fig. S9: Thermal performance curves 9
Fig. S10: Example of deterministic and stochastic population trajectories 10
Fig. S11: Longevity of co-occurrence as per the deterministic model 11
Fig. S12: Mean percent difference in co-occurrence relative to the deterministic model 12
Fig. S13: Fully factorial representation of co-occurrence 13
Fig. S14: Distributions of outcomes across select disturbance regimes 14
Fig. S15: Distribution of periods of co-occurrence by day 15
Fig. S16: Impact of disturbance size on co-occurrence by introduction day 16
Fig. S17: Impact of disturbance frequency on co-occurrence by introduction day 16
Fig. S18: Relationship between disturbance size and co-occurrence 17
Fig. S19: Extended analysis: thermal performance curves 18
Fig. S20: Extended analysis: summary plots 18
Fig. S21: Relationship between disturbance frequency and co-occurrence 19
Fig. S22: Distributions of co-occurrence and competitive outcomes 19
Fig. S23: Competitive outcomes by introduction day 20
Fig. S24: Fully factorial representation of transmission potential 21

**Sensitivity analysis**

We conducted a sensitivity analysis to determine how variations in (1) the value of $k$ and (2) the shape of the relationship between parasite replication rate and the temperature of the system would affect our results, as follows:

| | Original TPCs | Narrower TPCs | Wider TPCs |
|---|---|---|---|
| Original value of $k$ | OO | "ON" | "OW" |
| Higher value of $k$ | "HO" | "HN" | "HW" |
| Lower value of $k$ | "LO" | "LN" | "LW" |

Table S1: Table showing treatments used in sensitivity analysis and their corresponding abbreviations.

The higher value of $k$ was an order of magnitude larger than the original value of $k$. The lower value of $k$ was an order of magnitude smaller than the original value of $k$. Narrower TPCs increased maximum parasite replication rates, while wider TPCs decreased maximum parasite replication rates. Simulations were conducted using the $S_{DE}$ model; parasite replication rates peaked on either side of the average temperature of the system (offset responses scenario).

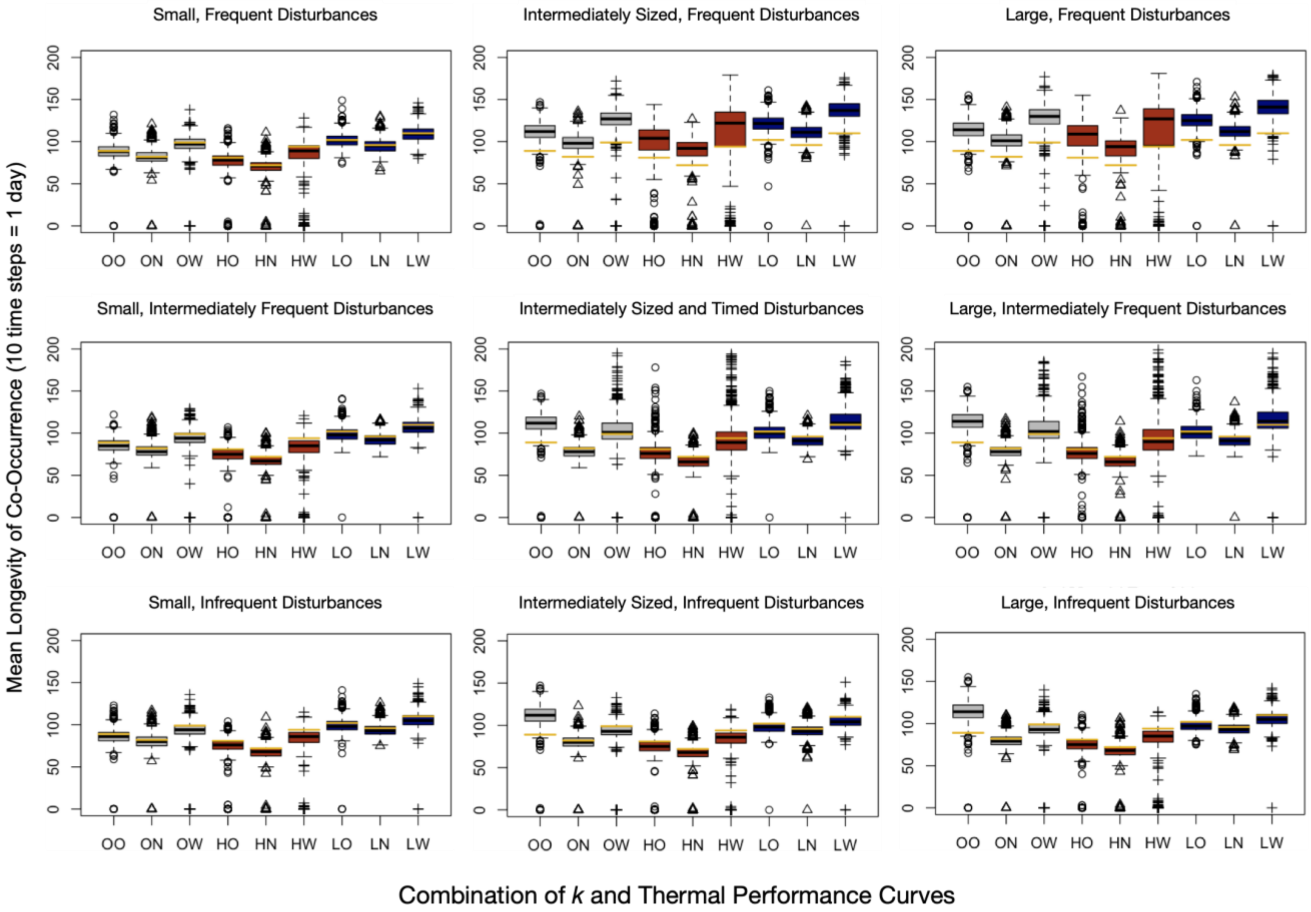


Figure S1: Simulations showing longevities of co-occurrence by treatment. Each panel shows the results of $S_{DE}$ model simulations under a given disturbance regime. Grey, red, and blue boxplots show the results of model simulations using the original, higher, and lower values of $k$. Circular, triangular, and crossed points show the results of model simulations within which replication rates are determined using the original, narrower, and wider thermal performance curves. Yellow lines denote deterministically simulated longevities of co-occurrence.

We found that all treatments produced qualitatively similar results to those presented in the main text.

**Extended analysis**

Using the Sharpe-Schoolfield thermal performance curves, we observed an initially positive and eventually saturating relationship between the magnitude of thermal disturbance and the longevity of co-occurrence. Given that the Sharpe-Schoolfield sub-model, cannot generate negative replication rates, extreme temperatures (which are more common when disturbances are large) while resulting in low replication rates, can never result in replication rates below zero, and thus, cannot drive down parasite populations. To investigate this response more thoroughly, we developed a set of quadratic thermal performance curves that could generate negative replication rates, and re-ran a subset of our simulations (Fig. S13). We found that when extreme temperatures result in replication rates below zero, the relationship between the magnitude of thermal disturbance and the longevity of co-occurrence is parabolic (Fig. S14).

Case 1: $P_i$ and $P_j$ respond identically to the temperature of the system

(1) $r_i = -0.005(t)^2 + 0.2(t)$

(2) $r_j = -0.005(t)^2 + 0.2(t)$

Where the temperature of the system ranges between -10°C and 50°C, with an average of 20°C.

Case 2: $P_i$ and $P_j$ respond identically to the temperature of the system

(1) $r_i = -0.005(t)^2 + 0.2(t)$

(2) $r_j = -0.005(t)^2 + 0.3(t) - 2.5$

Where the temperature of the system ranges between 0°C and 50°C, with an average of 25°C.

Case 3: $P_i$ and $P_j$ respond identically to the temperature of the system

(1) $r_i = -0.005(t)^2 + 0.2(t)$

(2) $r_j = -0.005(t)^2 + 0.3(t) - 2.5$

Where the temperature of the system ranges between 0°C and 50°C, with an average of 20°C.

Case 4: $P_i$ and $P_j$ respond identically to the temperature of the system

(1) $r_i = -0.005(t)^2 + 0.2(t)$

(2) $r_j = -0.005(t)^2 + 0.3(t) - 2.5$

Where the temperature of the system ranges between 0°C and 50°C, with an average of 30°C.

**High-average temperature environment analysis**
To investigate the effects of wider thermal performance curves (and thus wider thermal ranges), we developed a second set of thermal performance curves defined by a minimum temperature of 0°C, mean temperatures between 25.5 and 28°C, and a maximum temperature of 40°C. Within this environment, we re-ran all simulations described in the main text, but increased the range of disturbance size from 15 to 20 permutations. In summary, the outcomes of these simulations (detailed below) were qualitatively identical to those presented in the main text.

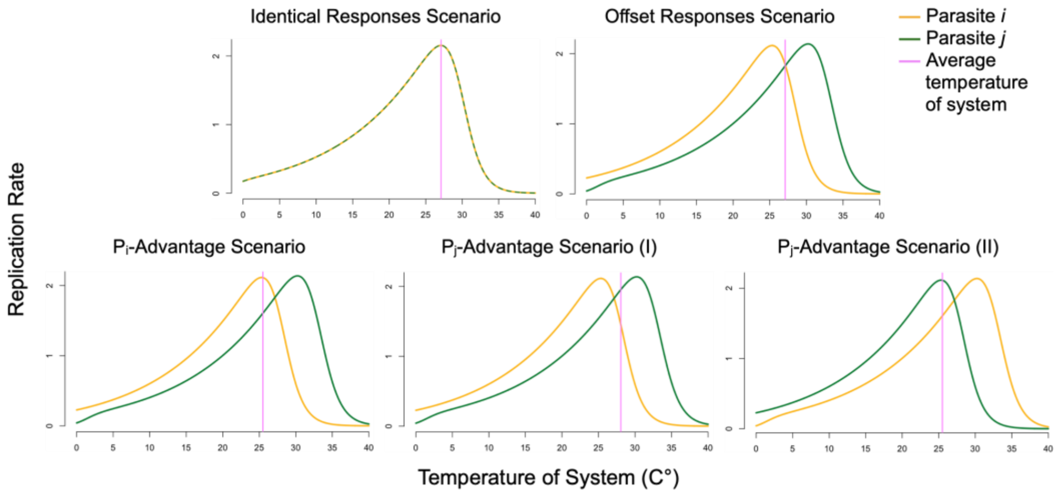


Figure S2: Sharpe-Schoolfield thermal performance curves relating the temperature of the system to the replication rates ($r_i$ and $r_j$) of two parasites ($P_i$ and $P_j$) in the High-Average Temperature Environment. Each panel (A-E) shows a different thermal scenario.

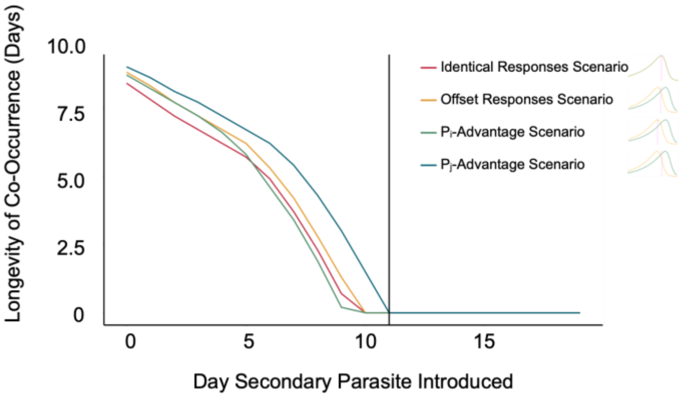


Figure S3: Deterministic longevities of co-occurrence in the High-Average Temperature Environment. The plot shows the number of time steps for which $P_i$ and $P_j$ have population abundances greater than one, by introduction day, as per the deterministic model. The black line denotes the first day on which the introduction of the secondary parasite, $P_j$, does not result in any co-occurrence between the two parasites.

When both parasites were introduced simultaneously, simulations of the demographically stochastic model ($S_D$) resulted in periods of co-occurrence that were, on average, 4.4% to 3.3% shorter (0.4 to 0.3 days) than deterministically simulated periods of co-occurrence (Figs. S4). Under the same conditions, simulations of the environmentally stochastic models ($S_E$, $S_{DE}$) resulted in periods of co-occurrence that were up to 62.1% ($S_E$) and 49.6% ($S_{DE}$) longer (5.2 ($S_E$) and 4.2 ($S_{DE}$) days) than deterministically simulated periods of co-occurrence. Across all 260 disturbance regimes, simulations of the $S_E$ model resulted in mean periods of co-occurrence that were between 3.4% shorter (0.3 days) and 62.1% longer (5.2 days) than their deterministic counterparts (Fig. S4). The identical responses scenario promoted the shortest and longest mean periods of co-occurrence (Figs. S4-S5). Simulations of the $S_{DE}$ model resulted in mean periods of co-occurrence that were between 6.8% shorter (0.6 days) and 49.6% longer (4.2 days) than deterministically simulated periods of co-occurrence (Figs. S4). Again, the identical responses scenario promoted the shortest and longest mean periods of co-occurrence (Figs. S4-S5). Across thermal scenarios, mean periods of co-occurrence were shortest when disturbances were large (standard deviation of the normal random variable in the temperature autocorrelation function = 20 to 40) and frequent (once every 6 to 9 days) (Figs. S4-S5). Mean periods of co-occurrence were longest when disturbances were large (SD = 34 to 40) and infrequent (five to ten times per day) (Fig. S5).

More generally, as the disturbances increased in size, periods of co-occurrence increased in length towards a saturation point (standard deviation of the normal random variable in the temperature autocorrelation function ≈ 22-40, depending on thermal scenario) (Fig. S6A, Supplementary Datafiles 3-1:5, 4-1:5). Additionally, when disturbances were infrequent (once every seven to ten days), periods of co-occurrence were similar in length to deterministically simulated periods of co-occurrence. Per the deterministic model, $P_i$ and $P_j$ can only co-occur for a maximum of nine days. Thus, systems subjected to infrequent disturbances were functionally similar to systems subjected to a constant environment, making the outcome of a simulation reliant on the initial conditions of the system. Interestingly, when disturbances were intermediately frequent (once per day to once every few days), periods of co-occurrence were most often shorter than deterministically simulated periods of co-occurrence (Fig. 6B). When disturbances were frequent (twice to ten times per day), mean periods of co-occurrence were longer than deterministically simulated periods of co-occurrence.

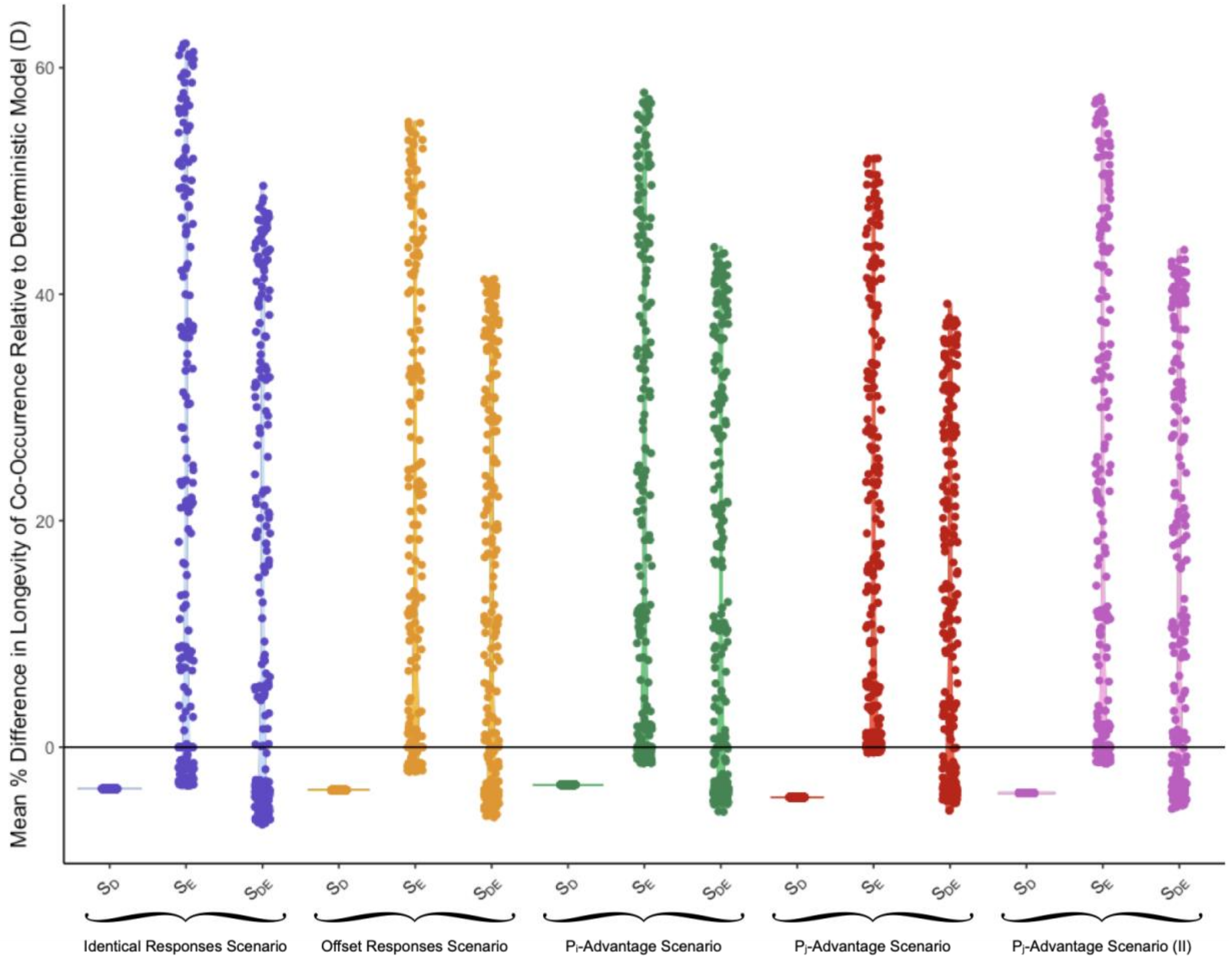


Figure S4: The mean percent difference in longevity of co-occurrence relative to deterministically simulated periods of co-occurrence in the High-Average Temperature Environment when both parasites are introduced at the same time. Each point represents the mean percent difference in longevity of co-occurrence relative to deterministically simulated periods of co-occurrence across 1000 stochastic simulations, as per a given environmental disturbance regime. As such, there are 260 points above each "$S_E$ Model"/"$S_{DE}$ Model" label.

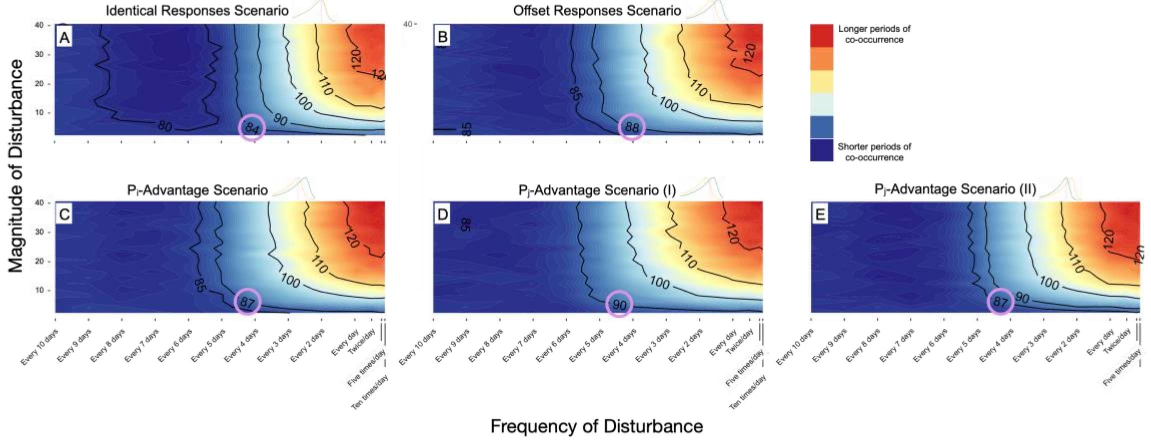


Figure S5: Fully factorial representations of the impacts of varyingly sized and timed disturbances on the longevity of co-occurrence in the High-Average Temperature Environment when both parasites are introduced at the same time. Each panel shows the mean number of time steps (10 time steps/day) for which $P_i$ and $P_j$ have population abundances greater than one as per the demographically and environmentally stochastic model, when $P_j$ has a thermal advantage. The circled numbers denote the number of time steps for which $P_i$ and $P_j$ can co-occur, per the deterministic model.

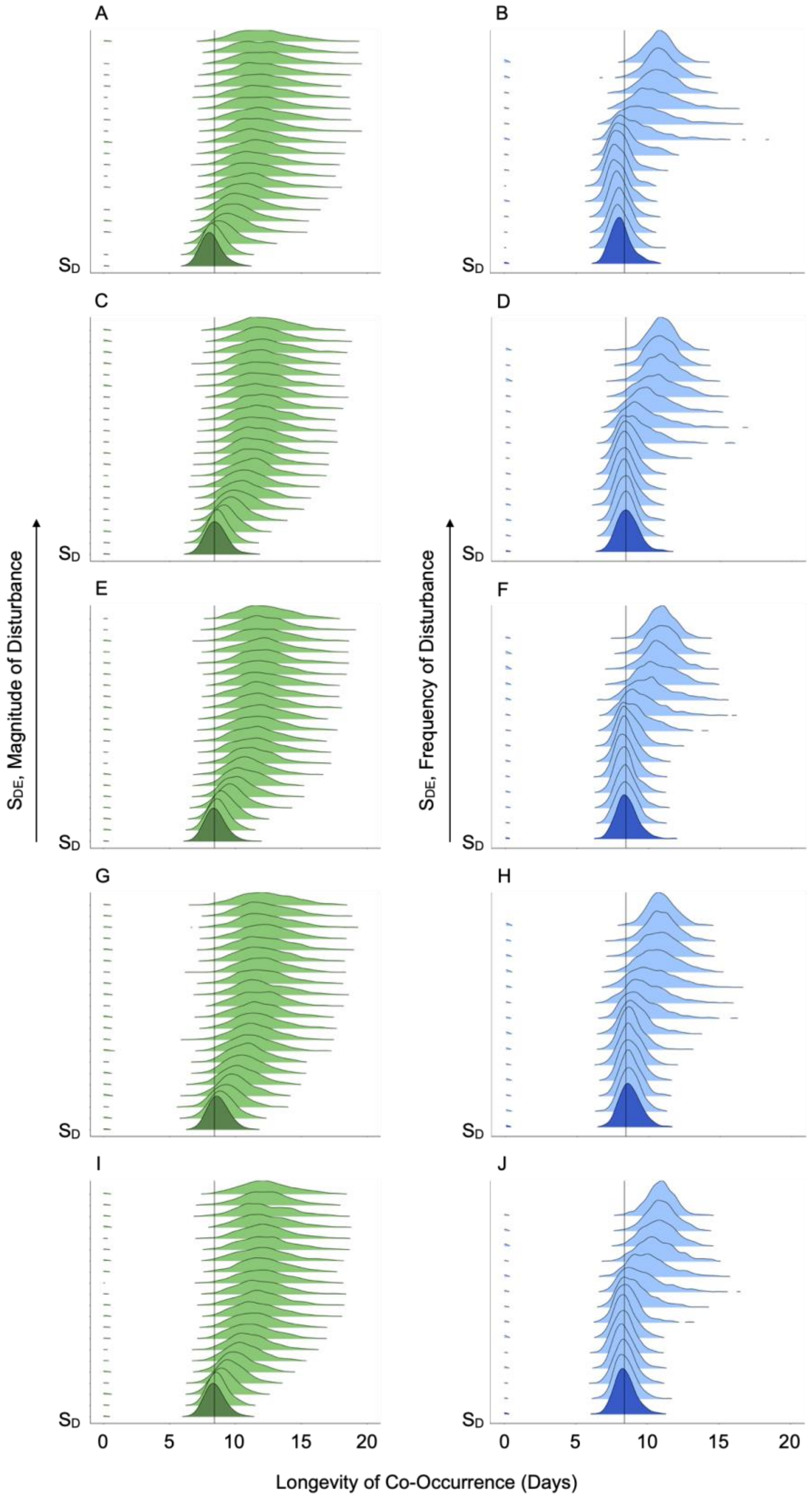


Figure S6: Distributions of outcomes across select disturbance regimes, as per the identical responses (A, B), offset responses (C, D), $P_i$-advantage (E, F), $P_i$-advantage (I) (G, H), and $P_i$-advantage (II) (I, J) scenarios in the High-Average Temperature Environment, when both parasites are introduced at the same time. "$S_D$" refers to longevities of co-occurrence as per the $S_D$ model. "$S_{DE}$" refers to longevities of co-occurrence as per the $S_{DE}$ model. Panels A, C, E, and G show the effects of increasingly large disturbances (frequency of disturbance fixed (once per day)). Panels B, D, F, and H show the effects of increasingly frequent disturbances (magnitude of disturbance fixed (SD = 10)). Black vertical lines represent deterministic reference points.

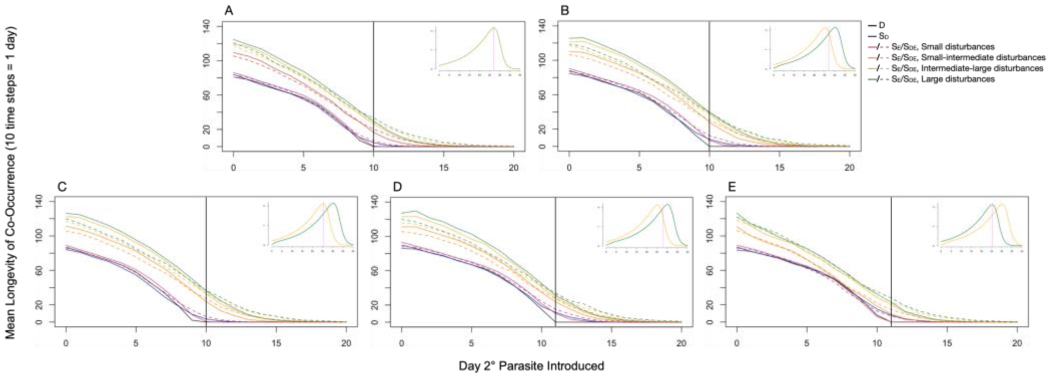


Figure S7: The impacts of varying magnitudes of disturbance on longevity of co-occurrence in the High-Average Temperature Environment. Each panel shows the mean number of time steps for which $P_i$ and $P_j$ have population abundances greater than one, by introduction day, when the magnitude of disturbance varies, and the frequency of disturbance is intermediate (once per day). Panels A-E show results from the identical responses, offset responses, $P_i$-advantage, and $P_j$-advantage (versions I and II) scenarios.

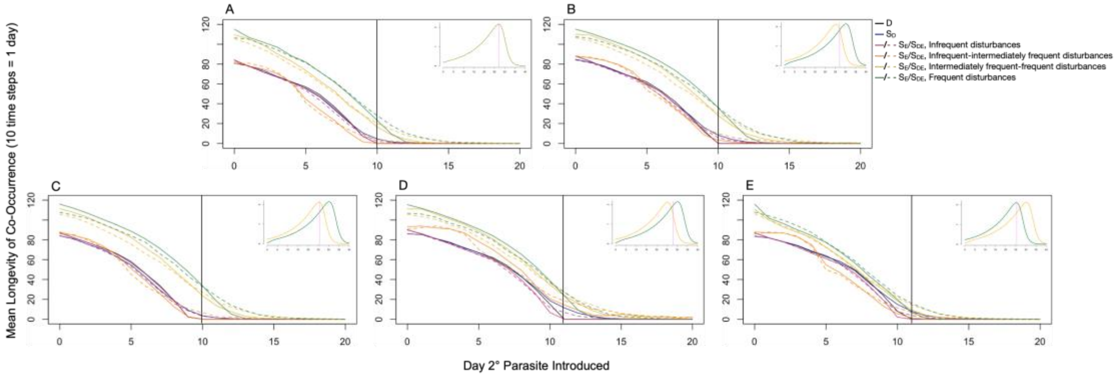


Figure S8: The impacts of varying frequencies of disturbance on longevity of co-occurrence in the High-Average Temperature Environment. Each panel shows the mean number of time steps for which $P_i$ and $P_j$ have population abundances greater than one, by introduction day, when the frequency of disturbance varies, and the magnitude of disturbance is intermediate (SD = 10). Panels A-E show results from the identical responses, offset responses, $P_i$-advantage, and $P_j$-advantage (versions I and II) scenarios.

### Sharpe-Schoolfield thermal sub-model

$$(1.1)\quad r_i(T(t+1)) = r_{i,0}e^{\frac{-E_{r_i}}{k}\left(\frac{1}{T(t+1)}-\frac{1}{T_0}\right)} \cdot \left(1 + e^{\frac{E^L_{r_i}}{k}\left(\frac{1}{T^L_{r_i}}-\frac{1}{T(t+1)}\right)} + e^{\frac{E^H_{r_i}}{k}\left(\frac{1}{T^H_{r_i}}-\frac{1}{T(t+1)}\right)}\right)^{-1}$$

$$(1.2)\quad r_j(T(t+1)) = r_{j,0}e^{\frac{-E_{r_j}}{k}\left(\frac{1}{T(t+1)}-\frac{1}{T_0}\right)} \cdot \left(1 + e^{\frac{E^L_{r_j}}{k}\left(\frac{1}{T^L_{r_j}}-\frac{1}{T(t+1)}\right)} + e^{\frac{E^H_{r_j}}{k}\left(\frac{1}{T^H_{r_j}}-\frac{1}{T(t+1)}\right)}\right)^{-1}$$

### Table S2: Parameter definitions and values for Sharpe-Schoolfield sub-model

| | | **Main Text Thermal Scenarios** | | | | | | | | **High-Average Temperature Environment Thermal Scenarios** | | | | | | | |
|---|---|---|---|---|---|---|---|---|---|---|---|---|---|---|---|---|---|
| | | **Identical** | | **Offset** | | **Pi-Advantage** | | **Pj-Advantage** | | **Identical** | | **Offset** | | **Pi-Advantage** | | **Pj-Advantage** | |
| | **Average Temperature of System** | 19.9+K | | 19.9+K | | 18+K | | 20.8530117+K | | 27.1+K | | 27.1+K | | 25.5+K | | 28.0639755+K | |
| **Symbol** | **Parameter** | **Pi Value** | **Pj Value** | **Pi Value** | **Pj Value** | **Pi Value** | **Pj Value** | **Pi Value** | **Pj Value** | **Pi Value** | **Pj Value** | **Pi Value** | **Pj Value** | **Pi Value** | **Pj Value** | **Pi Value** | **Pj Value** |
| $r_0$ | Baseline values of $r_i$ or $r_j$ at a reference temperature $T_0$ | 1.4 | 1.4 | 1.5 | 1.54 | 1.5 | 1.54 | 1.5 | 1.54 | 1.2 | 1.2 | 0.68 | 1.68 | 0.68 | 1.68 | 0.68 | 1.68 |
| $T_0$ (K) | Reference temperature (set arbitrarily such that $T^L$ << $T_0$ << $T^L$) | 14+K | 14+K | 13+K | 18+K | 13+K | 18+K | 13+K | 18+K | 19+K | 19+K | 11.3+K | 26+K | 11.3+K | 26+K | 11.3+K | 26+K |
| $E_r$ (eV) | Activation energy of $r_i$ or $r_j$ | 0.65 | 0.65 | 0.65 | 0.65 | 0.65 | 0.65 | 0.65 | 0.65 | 0.65 | 0.65 | 0.65 | 0.65 | 0.65 | 0.65 | 0.65 | 0.65 |
| $E_r^L$, $E_r^H$ (eV) | Low- and high-temperature inactivation energies of $r_i$ or $r_j$ | -6.2, 6.2 | -6.2, 6.2 | -6.2, 6.2 | -6.2, 6.2 | -6.2, 6.2 | -6.2, 6.2 | -6.2, 6.2 | -6.2, 6.2 | -6.2, 6.2 | -6.2, 6.2 | -6.2, 6.2 | -6.2, 6.2 | -6.2, 6.2 | -6.2, 6.2 | -6.2, 6.2 | -6.2, 6.2 |
| $T_r^L$, $T_r^H$ (K) | Low- and high-temperature inactivation thresholds for $r_i$ or $r_j$ | -9.5, 22.5+K | -9.5, 22.5+K | -12+K, 20.6+K | -7+K, 25.6K | -12+K, 20.6+K | -7+K, 25.6K | -12+K, 20.6+K | -7+K, 25.6K | -2+K, 29.8+K | -2+K, 29.8+K | -10+K, 28+K | 1+K, 33+K | -10+K, 28+K | 1+K, 33+K | -10+K, 28+K | 1+K, 33+K |
| k | Boltzmann's constant | - | - | - | - | - | - | - | - | - | - | - | - | - | - | - | - |

Where "K" = 273.15

**Fig. S9: Thermal performances curves**

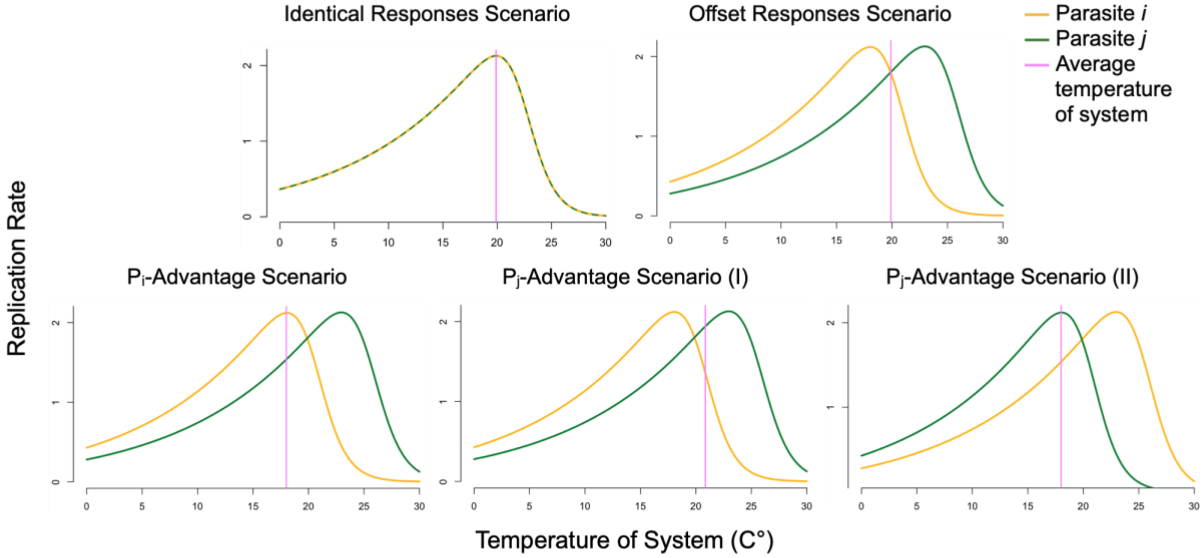


Figure S9: Sharpe-Schoolfield thermal performance curves relating the temperature of the system to the replication rates ($r_i$ and $r_j$) of two parasites ($P_i$ and $P_j$). Yellow lines show the relationship between the temperature of the system and the replication rate ($r_i$) of $P_i$. Green lines show the relationship between the temperature of the system and the replication rate ($r_j$) of $P_j$. Pink lines denote the average temperature of the system. Each panel (A-E) shows a different thermal scenarios.

**Fig. S10: Example of deterministic and stochastic population trajectories**

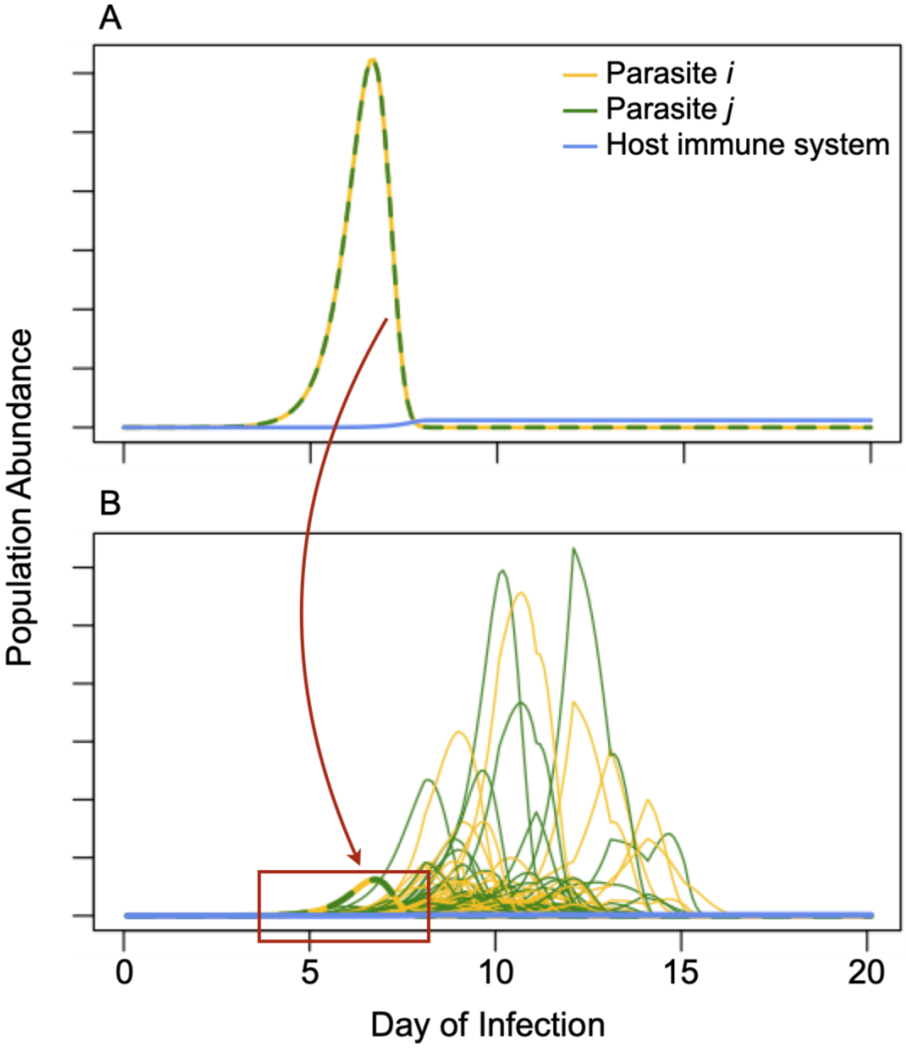


Figure S10: Example of deterministic and stochastic population trajectories. Panel A shows a simulation of the deterministic model. Parasite replication rates were given by the identical responses scenario, and the temperature of the system was held at the average temperature of the system (19.9°C). Panel B shows two hundred $S_{DE}$ model simulations. Parasite replication rates were given by the identical responses scenario, and thermal disturbances were intermediately sized and timed. In both cases, $P_i$ and $P_j$ were introduced simultaneously.

**Fig. S11: Longevity of co-occurrence as per the deterministic model**

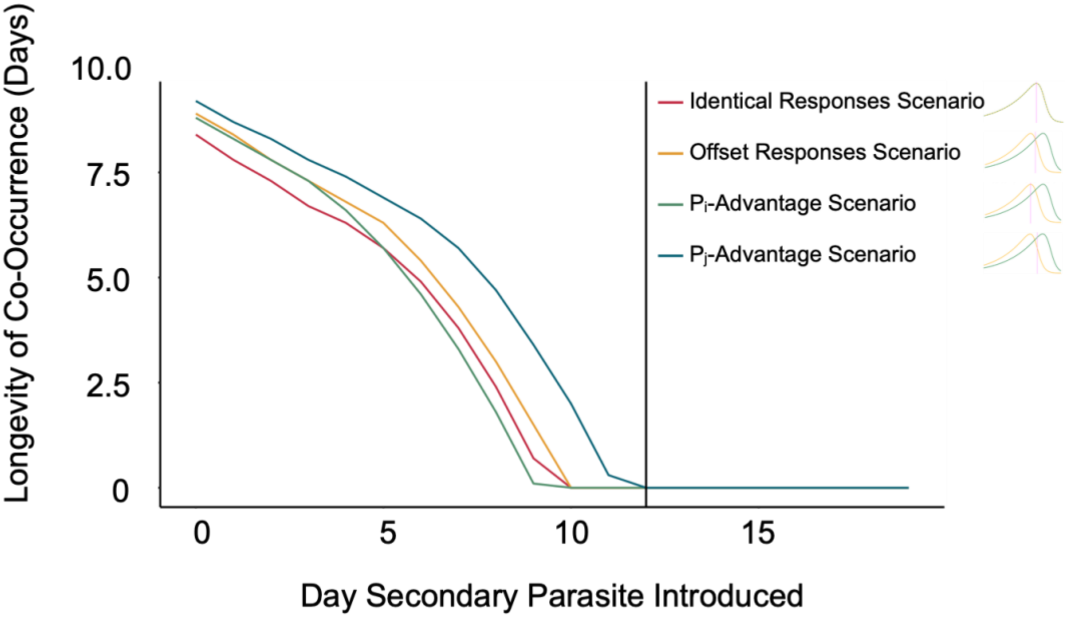


Figure S11: Longevity of co-occurrence as per the deterministic model. The plot shows the number of time steps for which $P_i$ and $P_j$ have population abundances greater than one, by introduction day, as per the deterministic model. The black line denotes the first day on which the introduction of the secondary parasite, $P_j$, does not result in any co-occurrence between the two parasites.

### Fig. S12: Mean percent difference in co-occurrence relative to the deterministic model

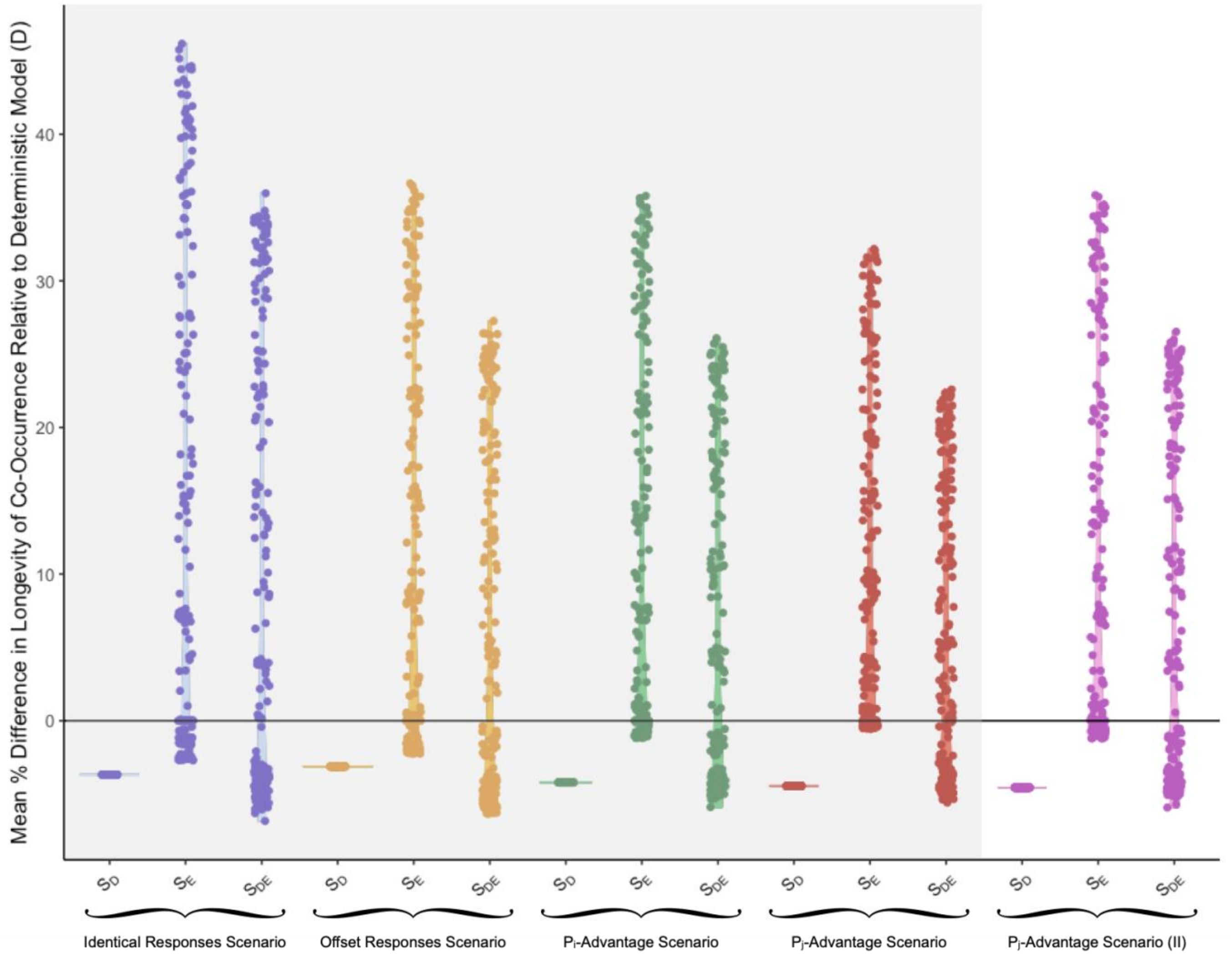


Figure S12: The mean percent difference in longevity of co-occurrence relative to the deterministic model, when both parasites are introduced at the same time. Each point represents the mean percent difference in longevity of co-occurrence relative to the deterministic model across 1000 stochastic simulations, as per a given environmental disturbance regime. As such, there is one point above each "$S_D$ Model" label, and 195 points above each "$S_E$ Model"/"$S_{DE}$ Model" label. This figure highlights results from the second version of the $P_j$-advantage scenario.

**Fig. S13: Fully factorial representation of co-occurrence**

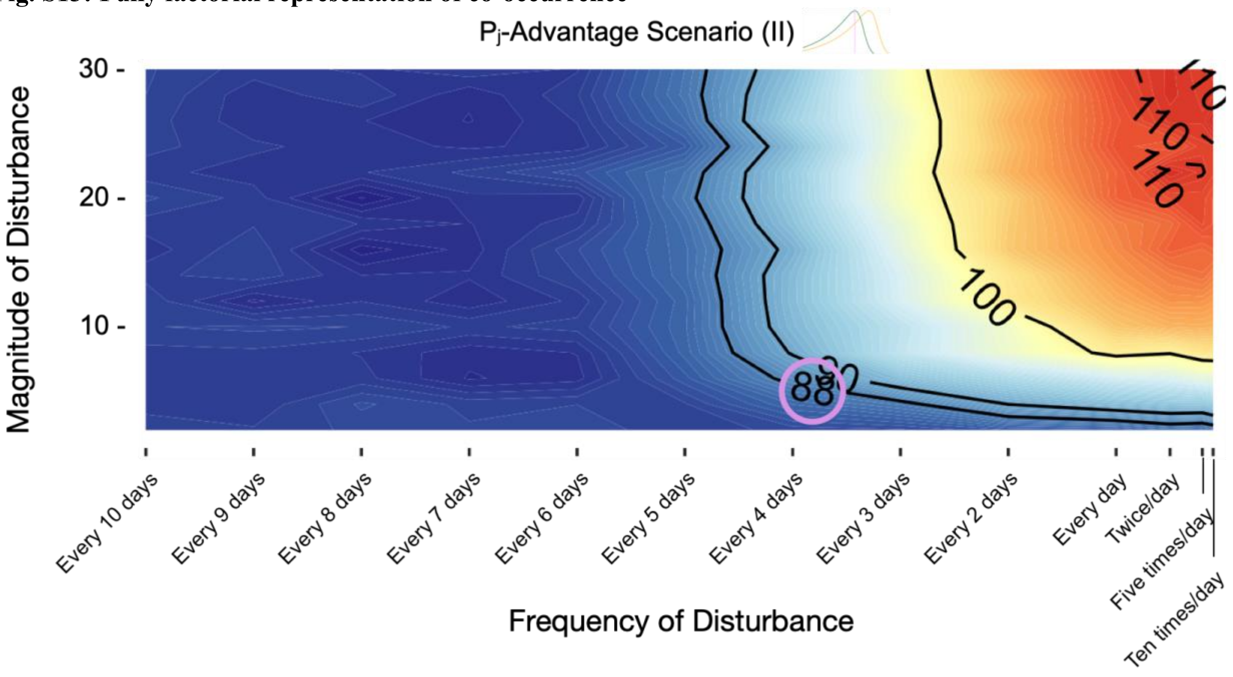


Figure S13: Fully factorial representations of the impacts of varyingly sized and timed disturbances on the longevity of co-occurrence when both parasites are introduced at the same time. The plot shows the mean number of time steps (10 time steps/day) for which $P_i$ and $P_j$ have population abundances greater than one as per the demographically and environmentally stochastic model, when $P_j$ has a thermal advantage. The circled number is the number of time steps for which $P_i$ and $P_j$ can co-occur as per the deterministic model.

**Fig. S14: Distributions of outcomes across select disturbance regimes**

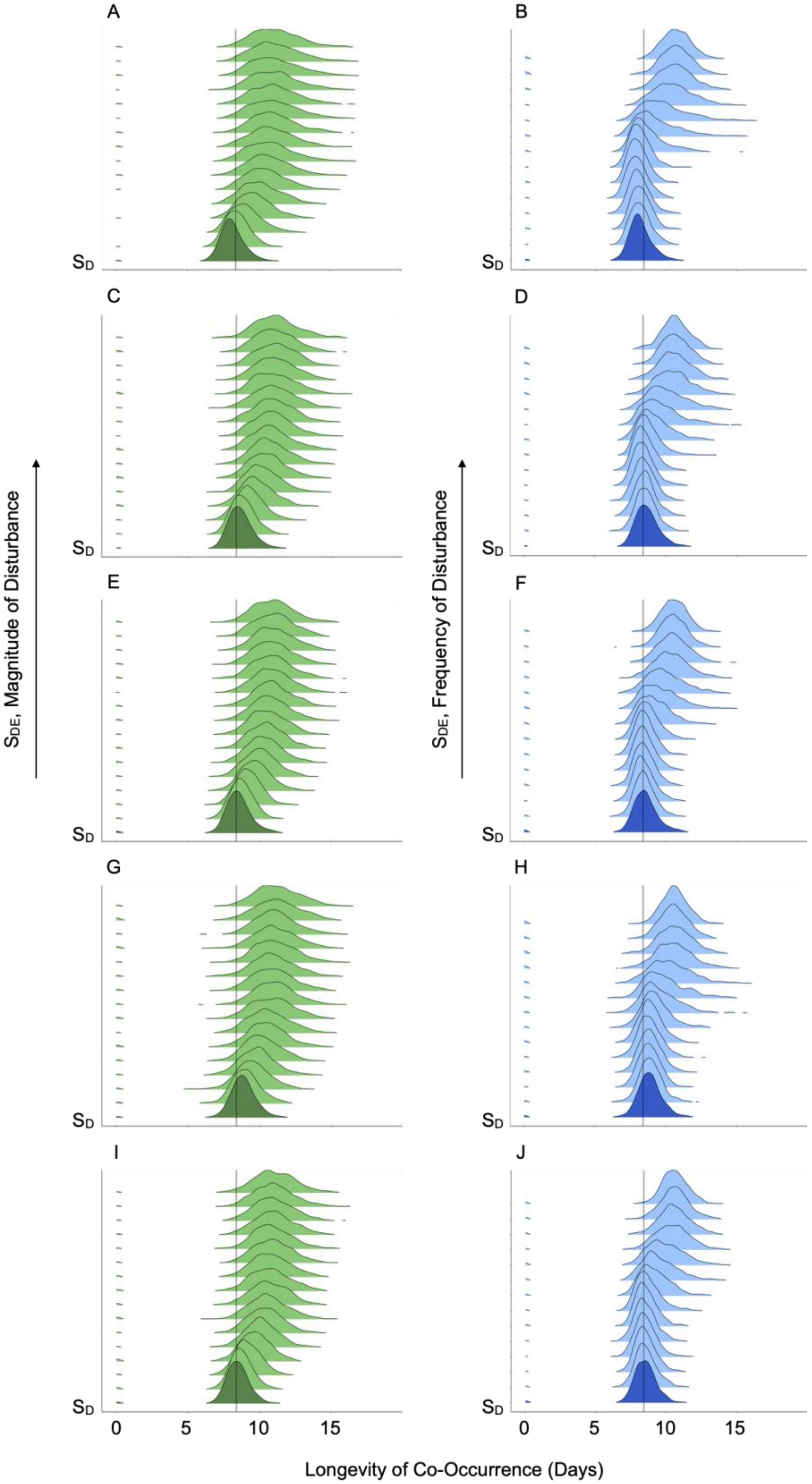


Figure S14: Distributions of outcomes across select disturbance regimes, as per the identical responses (A, B), offset responses (C, D), $P_i$-advantage (E, F), $P_j$-advantage (I) (G, H), and $P_j$-advantage (II) (I, J) scenarios, when both parasites are introduced at the same time. "$S_D$" refers to longevities of co-occurrence as per the $S_D$ model. "$S_{DE}$" refers to longevities of co-occurrence as per the $S_{DE}$ model. Panels A, C, E, G, and I show results from simulations in which disturbances increased in size, but were intermediately timed (once per day). Panels B, D, F, H, and J show results from simulations in which disturbances were intermediately sized (SD = 10), and increasingly frequent. The black vertical lines represent deterministic reference points.

**Fig. S15: Distribution of periods of co-occurrence by day**

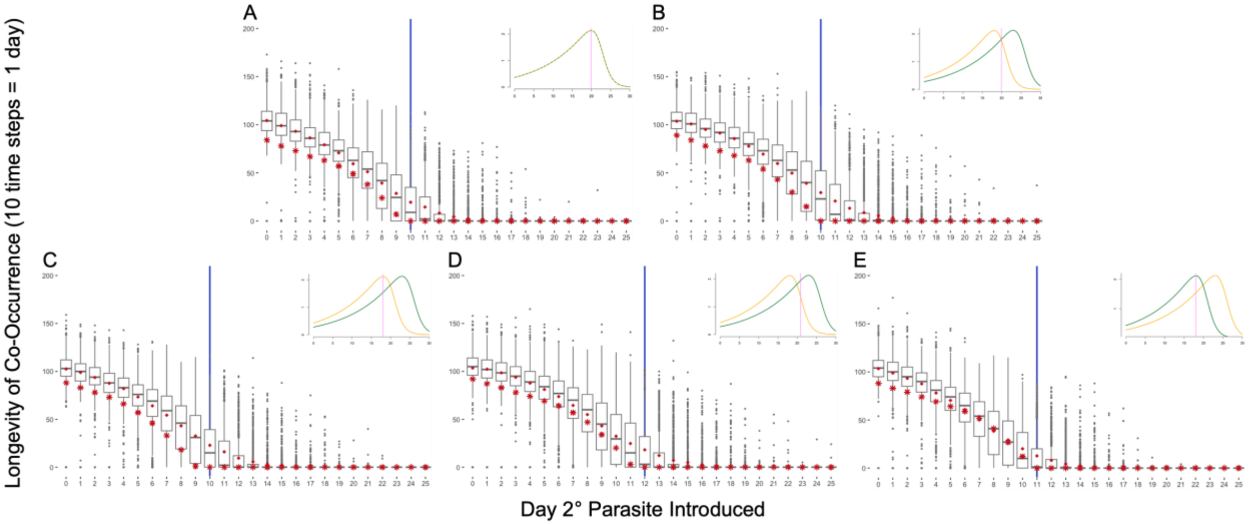


Figure S15: Simulation results. Each panel shows the number of time steps for which $P_i$ and $P_j$ have population abundances greater than one, by introduction day, as per the demographically and environmentally stochastic model. Panels A-E show results from the identical responses, offset responses, $P_i$-advantage, $P_j$-advantage (I), and $P_j$-advantage (II) scenarios when the disturbances are intermediately sized (SD = 10) and timed (once per day). Each dot shows the result of a single simulation. The grey lines represent medians, and red diamonds represent means. Red stars show the number of time steps for which $P_i$ and $P_j$ have population abundances greater than one as per the deterministic model. The blue line shows the first day on which the introduction of the secondary parasite, $P_j$, does not result in any co-occurrence between the two parasites.

### Fig. S16: Impact of disturbance size on co-occurrence by introduction day

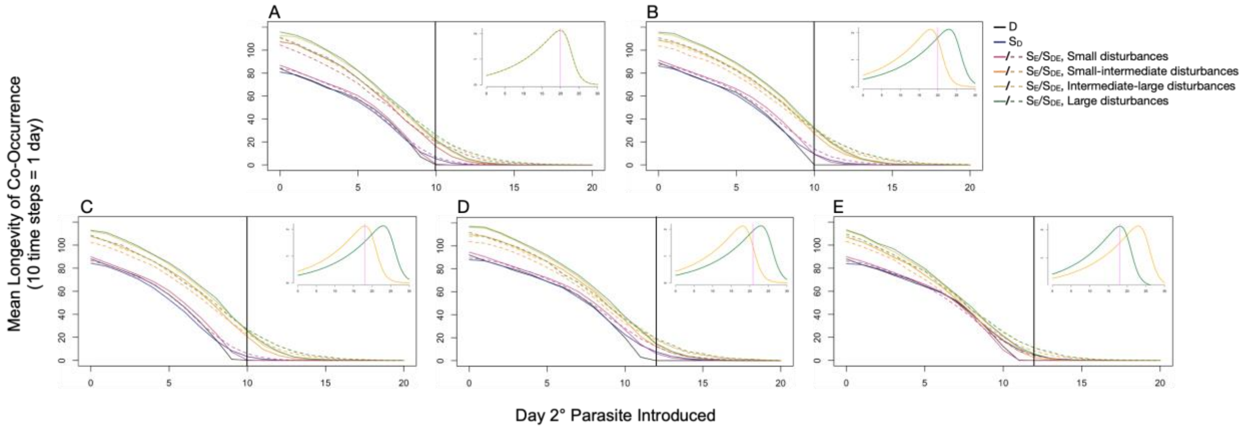


Figure S16: The impacts of varying magnitudes of disturbance on longevity of co-occurrence. Each panel shows the mean number of time steps for which $P_i$ and $P_j$ have population abundances greater than one, by introduction day, when the magnitude of disturbance varies, and the frequency of disturbance is intermediate (once per day). Panels A-E show results from the identical responses, offset responses, $P_i$-advantage, $P_j$-advantage (I), and $P_j$-advantage (II) scenarios.

### Fig. S17: Impact of disturbance frequency on co-occurrence by introduction day

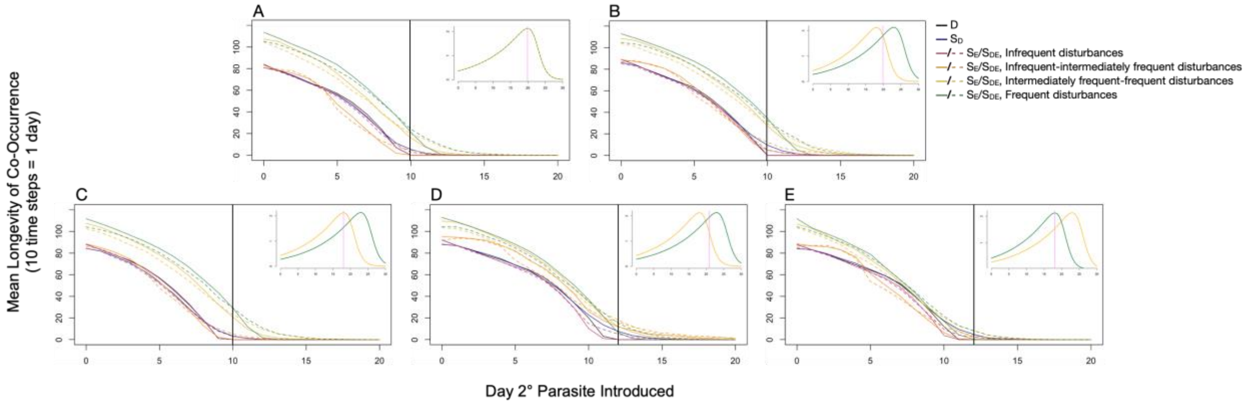


Figure S17: The impacts of varying frequencies of disturbance on longevity of co-occurrence. Each panel shows the mean number of time steps for which $P_i$ and $P_j$ have population abundances greater than one, by introduction day, when the frequency of disturbance varies, and the magnitude of disturbance is intermediate (SD = 10). Panels A-E show results from the identical responses, offset responses, $P_i$-advantage, $P_j$-advantage (I), and $P_j$-advantage (II) scenarios.

**Fig. S18: Relationship between disturbance size and co-occurrence**

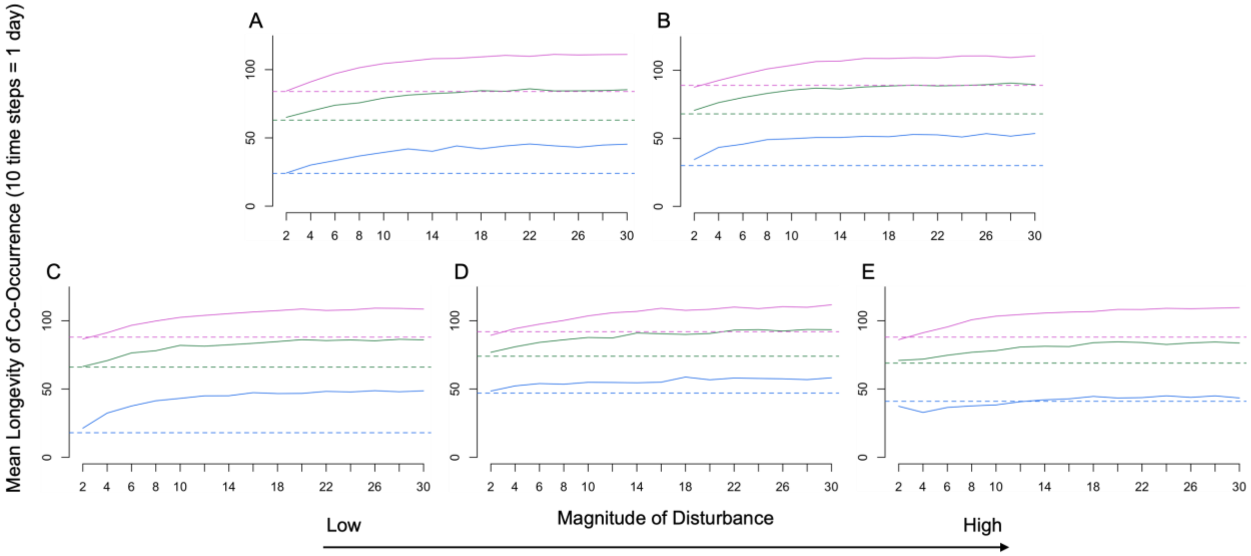


Figure S18: The impacts of varying magnitudes of disturbance on longevity of co-occurrence. Each panel shows the mean number of time steps for which $P_i$ and $P_j$ have population abundances greater than one by magnitude of disturbance when the frequency of disturbance is intermediate (once per day), as per the demographically and environmentally stochastic model. Solid and dashed lines show mean longevities and deterministic longevities, respectively, for comparison. Pink, green, and blue lines show outcomes resulting from the secondary parasite being introduced on Day 0, Day 4, and Day 8, respectively. Panels A-E show results from the identical responses, offset responses, $P_i$-advantage, $P_j$-advantage (I), and $P_j$-advantage (II) scenarios.

### Fig. S19: Extended analysis: thermal performance curves

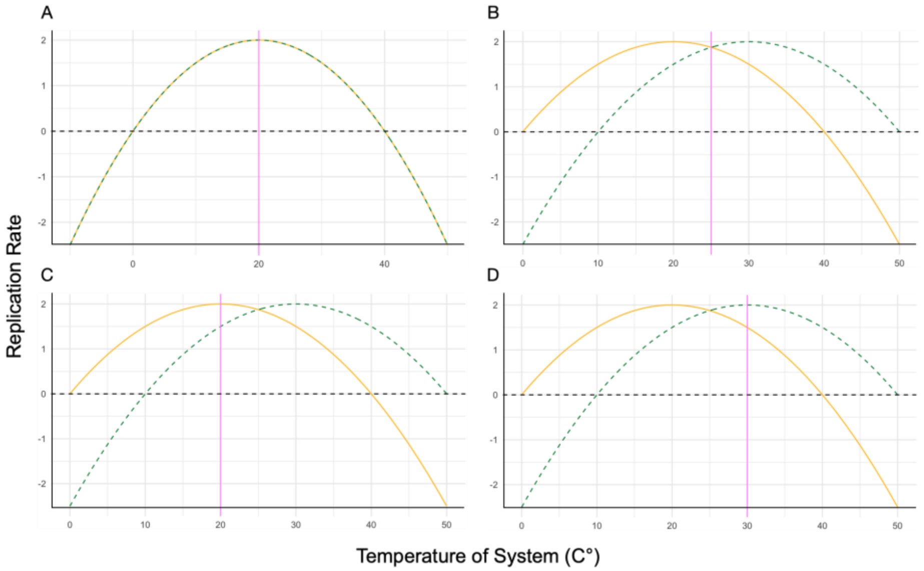


Figure S19: Quadratic thermal performance curves used to explore the implications of negative replication rates on the relationship between magnitude of disturbance and longevity of co-occurrence. Yellow lines show the relationship between the temperature of the system and the replication rate ($r_i$) of $P_i$. Green lines show the relationship between the temperature of the system and the replication rate ($r_j$) of $P_j$. Pink lines denote the average temperature of the system. Panels A-D show the identical responses, offset responses, $P_i$-advantage, $P_j$-advantage (I) scenarios.

### Fig. S20: Extended analysis: summary plots

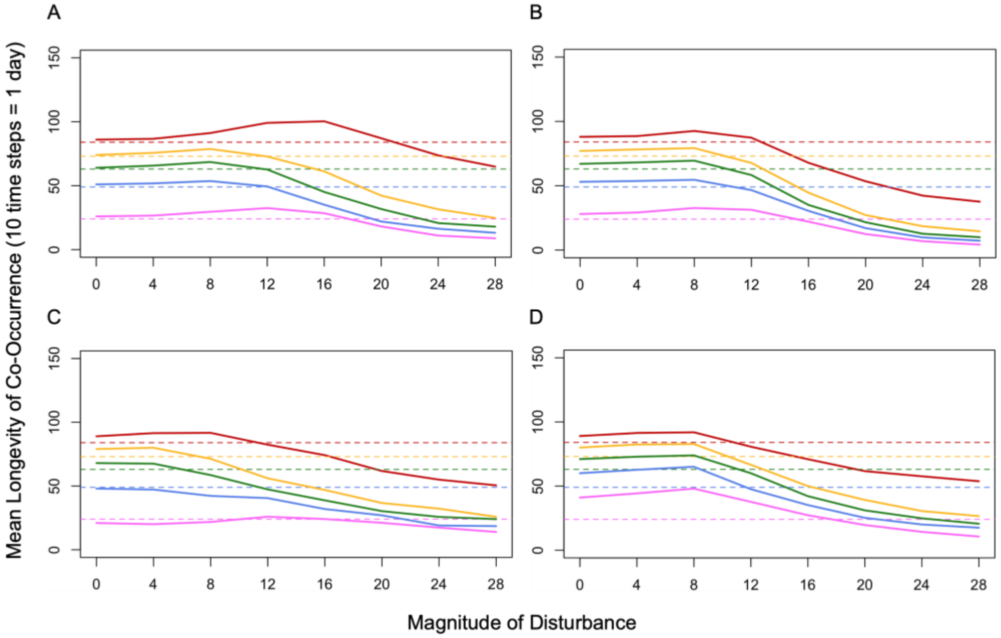


Figure S20: The impacts of varying magnitudes of disturbance on longevity of co-occurrence when extreme temperatures produce negative replication rates. Each panel shows the mean number of time steps for which $P_i$ and $P_j$ have population abundances greater than one by magnitude of disturbance when the frequency of disturbance is intermediate (once per day), as per the environmentally stochastic model. Solid and dashed lines show mean longevities and deterministic longevities, respectively, for comparison. Red, yellow, green, blue, and pink lines show outcomes resulting from the secondary parasite being introduced on Day 0, Day 2, Day 4, Day 6, and Day 8, respectively. Panels A-D show results from the identical responses, offset responses, $P_i$-advantage, $P_j$-advantage (I) scenarios.

**Fig. S21: Relationship between disturbance frequency and co-occurrence**

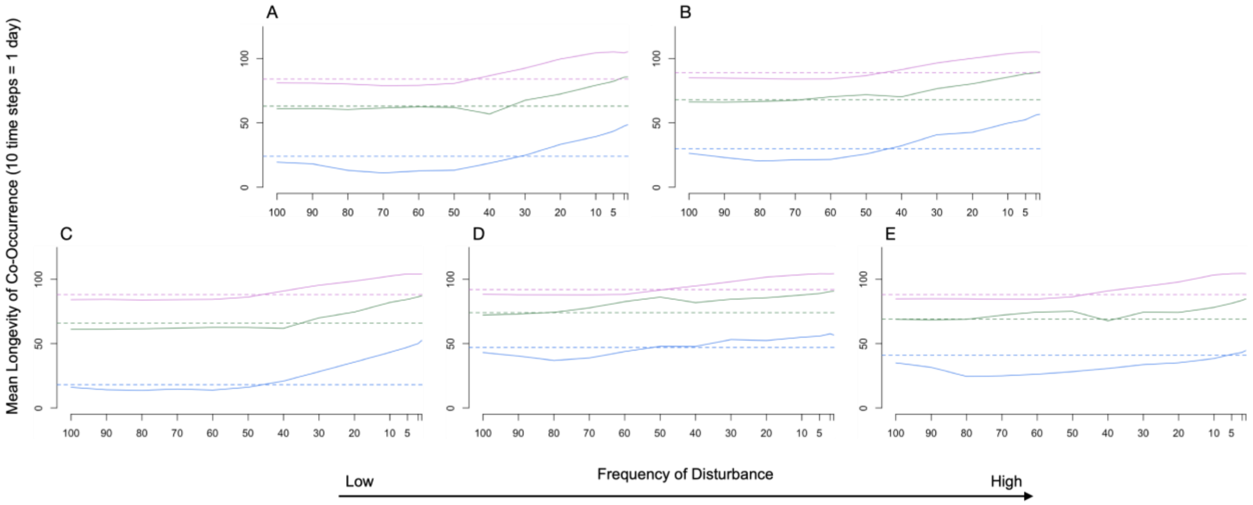


Figure S21: The impacts of varying frequencies of disturbance on longevity of co-occurrence. The plot shows the mean number of time steps (10 time steps/day) for which $P_i$ and $P_j$ have population abundances greater than one by frequency of disturbance when the magnitude of disturbance is intermediate (SD = 10), as per the demographically and environmentally stochastic model. Solid and dashed lines show mean longevities and deterministic longevities, respectively, for comparison. Pink, green, and blue lines show outcomes resulting from the secondary parasite being introduced on Day 0, Day 4, and Day 8, respectively. Panels A-E show results from the identical responses, offset responses, $P_i$-advantage, $P_j$-advantage (I), and $P_j$-advantage (II) scenarios.

### Fig. S22: Distributions of co-occurrence and competitive outcomes

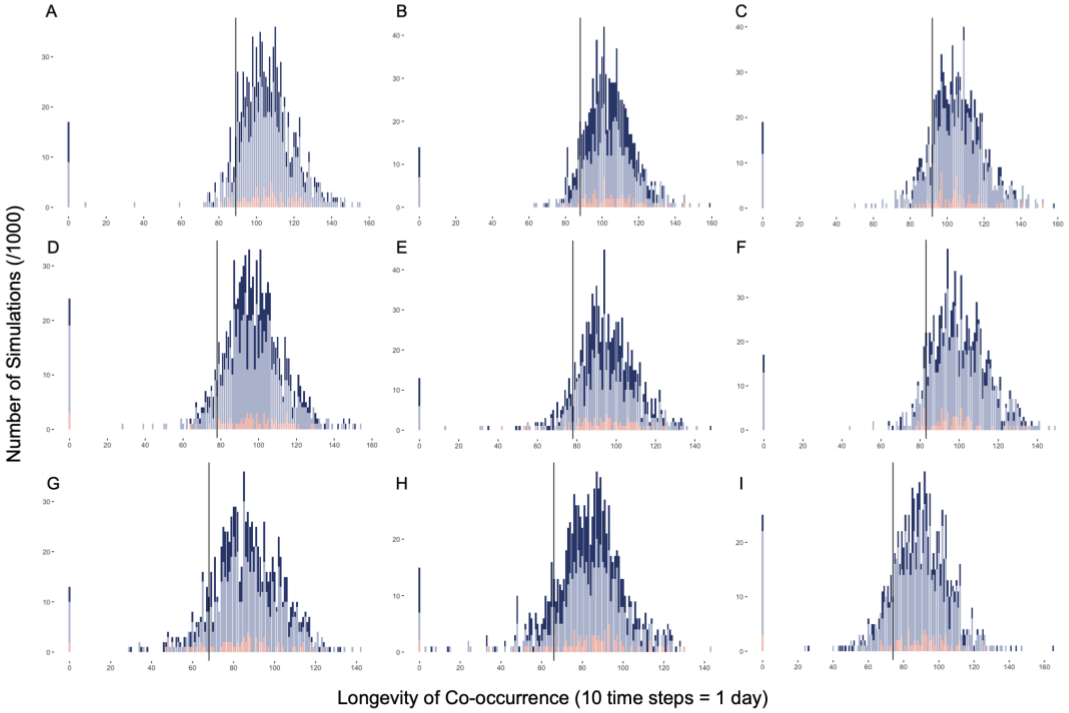


Figure S22: Distribution of outcomes, and outcomes of competition in each of 1000 simulations. Each panel shows the number of simulations, out of 1000, that resulted in a given period of co-occurrence, by competitive outcome (i.e. $P_i$ outlasts $P_j$ (dark blue), $P_j$ outlasts $P_i$ (light blue), and $P_i$ and $P_j$ are simultaneously extirpated (pink)), as per the demographically and environmentally stochastic model. Each row shows results from the offset responses, $P_i$-advantage, and $P_j$-advantage (I) scenarios when disturbances were intermediately sized (SD = 10) and timed (once per day). In the first, second, and third rows, the secondary parasite ($P_j$) was introduced on Day 0, Day 2, and Day 4, respectively.

### Fig. S23: Competitive outcomes by introduction day

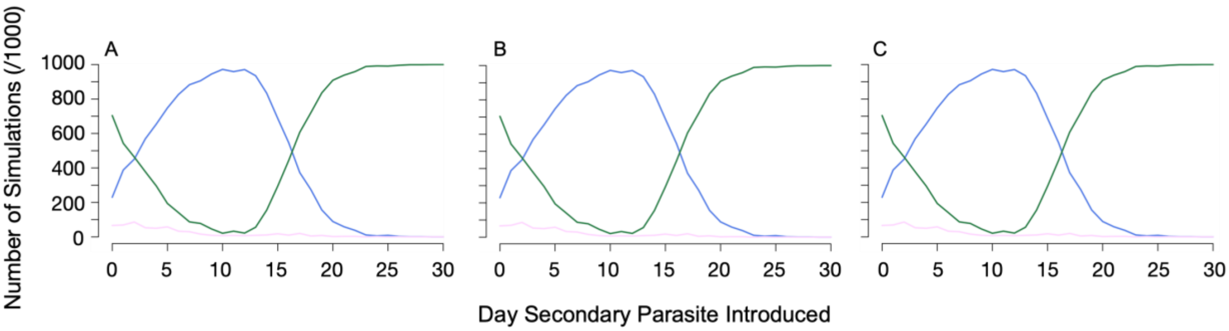


Figure S23: Summarised outcomes of competition in each of 1000 simulations. Each panel shows the number of simulations, out of 1000, that resulted in a given competitive outcome (i.e. $P_i$ outlasts $P_j$ (blue), $P_j$ outlasts $P_i$ (green), and $P_i$ and $P_j$ are simultaneously extirpated (pink)), by introduction time, as per the demographically and environmentally stochastic model. Panels A, B, and C show results from the offset responses, $P_i$-advantage, and $P_j$-advantage (I) scenarios when disturbances were intermediately sized (SD = 10) and timed (once per day).

**Fig. S24: Fully factorial representation of transmission potential**

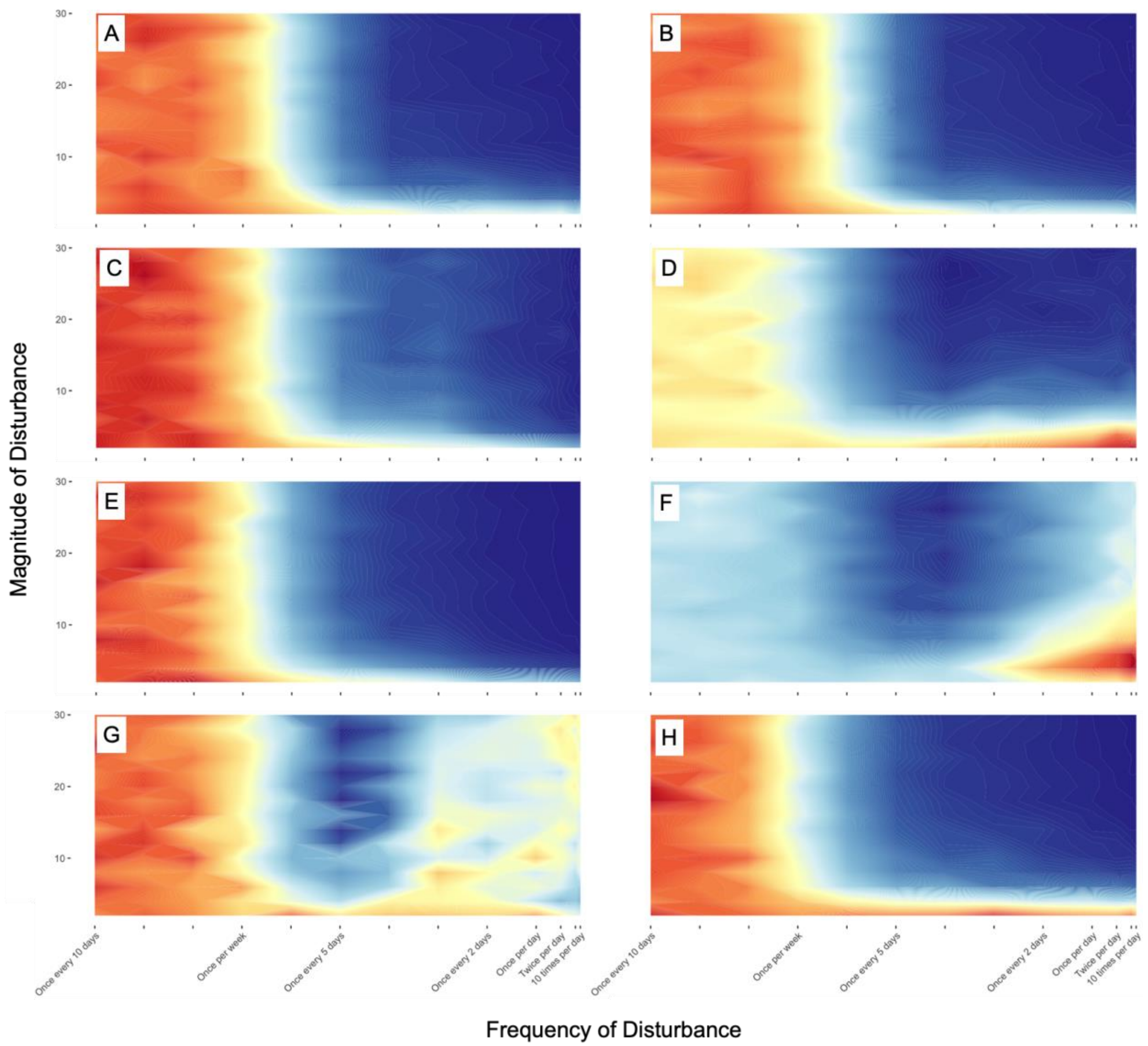


Figure S24: Fully factorial representations of the impacts of varyingly sized and timed disturbances on the transmission potentials (TPs) of co-occurring parasites. Each panel shows the median TP of co-co-occurring parasites as per the demographically and environmentally stochastic model. The first column shows median TPs of the primary parasite ($P_i$); the second column shows median TPs of the secondary parasite ($P_j$). The first, second, third, and fourth rows show results from the identical responses, offset responses, $P_i$-advantage, and $P_j$-advantage (I) scenarios. Warmer colours denote higher TPs.
